# Optimal Pre-Experimental Coil Sequence Selection for TMS Motor Mapping

**DOI:** 10.64898/2026.05.26.727891

**Authors:** Renxiang Qiu, Ole Numssen, Benjamin Kalloch, Florian Wilhelmy, Erdem Güresir, Konstantin Weise, Thomas R. Knösche

**Affiliations:** Max Planck Institute for Human Cognitive and Brain Sciences, Leipzig, Germany; Department of Neurosurgery, University Hospital Leipzig, Leipzig, Germany; Leipzig University of Applied Sciences (HTWK), Leipzig, Germany; Technische Universität Ilmenau, Ilmenau, Germany

**Keywords:** Transcranial Magnetic Stimulation, Motor Mapping and Optimal Experimental Design

## Abstract

Transcranial magnetic stimulation (TMS) motor mapping increasingly relies on electric-field (E-field) modeling to localize cortical targets, but many candidate coil placements induce highly redundant cortical patterns. We frame prospective coil-sequence design as a subset-selection problem and compare farthest-point sampling, determinant-based objectives, and related controls in virtual mapping experiments across 12 realistic head models. Across convergence, matrix-diagnostic, and manifold-coverage analyses, the best-performing objectives combined low between-stimulation redundancy with preserved inter-element separability on the cortex; objectives that maximized one of these at the expense of the other fell below random sampling. An RBF-kernelized D-optimal objective matched FPS in mapping accuracy while using ∼15 fewer unique scalp positions per 100 pulses, suggesting reduced arm-reconfiguration cost on robotic TMS platforms. A singular-spectrum analysis of the candidate library provides a low-cost a-priori indicator that predicts when discretization choices fall below the diversity floor required for stable mapping. These results recast prospective TMS mapping as a limited manifold-coverage problem and offer concrete design rules for sample-efficient, robotically efficient sequence planning. Algorithm implementations are available as part of the open-source `pynibs` tool-box at https://gitlab.gwdg.de/tms-localization/pynibs/-/tree/dev/pynibs/optimization.

## 1 Introduction

Transcranial magnetic stimulation (TMS) is a widely used non-invasive method for studying human brain function by inducing brief intracranial electric fields (E-fields) (Barker et al., 1985). In the motor system, single-pulse stimulation over the precentral gyrus can elicit motor evoked potentials (MEPs) in contralateral muscles, providing a physiological readout for identifying stimulation targets and characterizing individual motor representations. Precise motor maps are valuable in research and clinical practice, including presurgical planning and individualized targeting of therapeutic stimulation (Krieg et al., 2014; Rossini et al., 2015). Yet the effective cortical locus of a TMS pulse is not determined by scalp position alone: it also depends on coil orientation, coil-to-cortex distance, individual anatomy, and tissue conductivity, so scalp coordinates remain an imperfect proxy for the stimulated cortical site (Opitz et al., 2013; Thielscher et al., 2011; Vetter et al., 2025).

Traditional motor-mapping protocols typically sample a grid of scalp locations and summarize MEP amplitudes to estimate a scalp “hotspot”, which is then projected onto cortex. This workflow is straightforward to implement, but it largely ignores the spatial distribution of the induced E-field. Subject-specific finite-element simulations offer a more direct route to cortical localization by estimating the field associated with each stimulation, and E-field-aware mapping approaches use these simulations to infer candidate cortical sites more directly than scalp-based proxies (Aonuma et al., 2018; Opitz et al., 2013; Weise et al., 2020).

Recent E-field-informed statistical mapping methods relate local cortical E-field strength to MEP amplitudes across many stimulations, fit element-wise response curves, and summarize fit quality as goodness-of-fit (e.g., *R*^2^) maps that identify candidate cortical origins of the MEP (Numssen et al., 2021; Weise et al., 2020, 2023). The main opportunity for improvement is sampling efficiency: reliable inference requires coil placements that generate distinct E-field patterns within the ROI, and the problem becomes still harder outside motor cortex, where higher response variability weakens the local E-field–readout relation (Bhutto et al., 2026; Jing et al., 2023). Random or pseudo-random sampling has reduced experimental burden relative to earlier multi-condition designs (Numssen et al., 2021; Weise et al., 2023), but does not explicitly prioritize informative placements; because TMS-induced fields vary smoothly across nearby coil poses, neighboring placements yield highly correlated cortical patterns (Schultheiss et al., 2025; Thielscher et al., 2011; Weise et al., 2020).

These constraints motivate *prospective* coil-sequence design: selecting coil placements from predicted E-field characteristics in advance of the TMS session. Schultheiss et al. proposed a prospective sequence-design strategy based on farthest point sampling (FPS), which constructs a sequence by maximizing Euclidean separation between E-field patterns (Schultheiss et al., 2025). More generally, prospective coil-sequence selection can be cast as a design-of-experiments problem over a discrete library of candidate E-field patterns, with objective functions that emphasize different notions of redundancy, coverage, and information.

Optimal experimental design provides one useful perspective. When measurements are costly, determinant-based (D-optimal) criteria seek designs that reduce uncertainty in the estimated coefficients of a linear measurement model by maximizing the determinant of an information matrix (Joshi & Boyd, 2009). Related determinant-based objectives have also been used for sensor selection and admit efficient greedy implementations (Manohar et al., 2018; Saito et al., 2021). In the TMS setting, however, the behavior of such objectives depends on the geometry and effective dimensionality of the feasible E-field library, which have not been systematically characterized for this sequence-design problem. We therefore explicitly examine how candidate E-field patterns populate the attainable stimulation manifold and how different objectives traverse it.

Prospective design also depends on how the candidate library is discretized. Schultheiss et al. (Schultheiss et al., 2025) showed empirically, by running full FPS-based mapping at every setting, that spatial and angular step saturate quickly whereas search radius materially changes mapping behavior. We recast this as a library-geometry question and use the singular spectrum of the candidate E-field library as an algorithm-agnostic diagnostic of which discretization choices actually broaden the attainable E-field space, with a cross-algorithm robustness check for the radius axis.

Here we present a unified framework for prospective coil-sequence selection in E-field-informed motor mapping. We compare methods along three design choices: the pattern representation, the redundancy metric, and the set-level aggregation rule. Because the resulting sequences are computed offline and can be rearranged at execution time (e.g., by grouping stimulations at the same scalp position), the choice of objective affects movement cost on robotic or neuronavigated TMS systems as well as statistical efficiency. Using anatomically realistic head models and virtual mapping simulations, we quantify how these choices affect redundancy, map convergence, and coil-pose usage; relate the differences to the geometry of the feasible E-field manifold; and derive practical guidance for sequence planning, including settings without robotic positioning (Gomez-Tames et al., 2018).

## 2 Methods

### 2.1 Problem formulation and the coil selection challenge

Prospective TMS sequence design is a continuous planning problem over scalp positions, coil orientations, coil-to-cortex distance, and safety constraints. We discretize this space into a finite candidate library so that E-field simulations and combinatorial selection are tractable, and use greedy algorithms for the latter.

Let *C* = {1, …, *N* } index a discrete library of candidate TMS coil placements (positions and orientations) distributed over the scalp above the target cortical region of interest (ROI). For each candidate placement *i* ∈ *C*, we pre-compute the induced E-field magnitude at each ROI element via high-resolution finite-element simulation (Section 2.2). Stacking these patterns yields an *E-field library matrix* **E** ∈ ℝ^*N* ×*V*^, where *N* is the number of candidate coil placements, *V* is the number of cortical elements in the ROI, and each row **e**_*i*_ ∈ ℝ^*V*^ represents the E-field pattern induced by placement *i*.

Our goal is to select an optimal subset of *K* coil placements from *C*. The combinatorial target is a set (Eq. 1), but greedy selection returns an ordered sequence *S* = (*s*_1_, …, *s*_*K*_) whose prefix *S*_*k*_ = (*s*_1_, …, *s*_*k*_) is itself the greedy solution for budget *k*; this prefix property is exploited in the convergence analyses. We use “sequence” for the ordered greedy output and “subset” for the unordered target. The optimization problem is

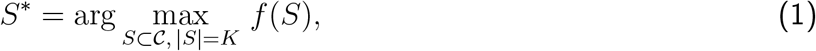

where *f* (*S*) quantifies the diversity or information content of the selected set. The central question is which objective *f* (·) best captures the informational value of a coil-placement set for accurate motor mapping, measured by hotspot localization error (geodesic distance) and map fidelity (NRMSD), under a limited pulse budget.

### 2.2 Candidate library and E-field simulation

To construct subject- and ROI-specific E-field libraries, we built high-resolution finite-element head models from structural MRI data. T1-and T2-weighted scans were segmented and meshed with the SimNIBS 4.5 `charm` pipeline (Puonti et al., 2020). The models comprised nine tissue classes with conductivity values taken from the SimNIBS configuration (Appendix A) (Puonti et al., 2020; Saturnino et al., 2019). Diffusion-weighted images were used to reconstruct anisotropic conductivity profiles in gray and white matter via volume-normalized tensor mapping (Güllmar et al., 2010; Opitz et al., 2011).

The ROI covered dorsal somatosensory cortex (BA 1 and BA 3), primary motor cortex (BA 4), and premotor cortex (BA 6). We created the mask on the FreeSurfer average template and transformed it to each individual head model. The figure-8 D70 Air-Film coil (Magstim Ltd., Whitland, UK) was modeled with 10,000 magnetic dipoles arranged in five layers in the coil plane, with dipole magnitudes derived from the enclosed current (Weise et al., 2023).

All analyses were performed on the mid-layer of the gray-matter compartment to reduce boundary effects caused by conductivity discontinuities. E-field magnitudes at the mid-layer were obtained with superconvergent patch recovery (Saturnino et al., 2019; Zienkiewicz & Zhu, 1992).

#### Candidate coil placements

We sampled candidate coil center positions uniformly on the scalp surface overlying the ROI (437 positions, ∼2.5 mm spacing) and, at each position, simulated 12 tangential coil orientations at 15°intervals over the half-circle 0°–165°relative to the posterior–anterior axis, giving *N* = 437 × 12 = 5,244 candidate placements per subject (Figure 1). Discretization sensitivity is characterized using the singular-spectrum analysis defined in Section 2.4.

**Figure 1:**
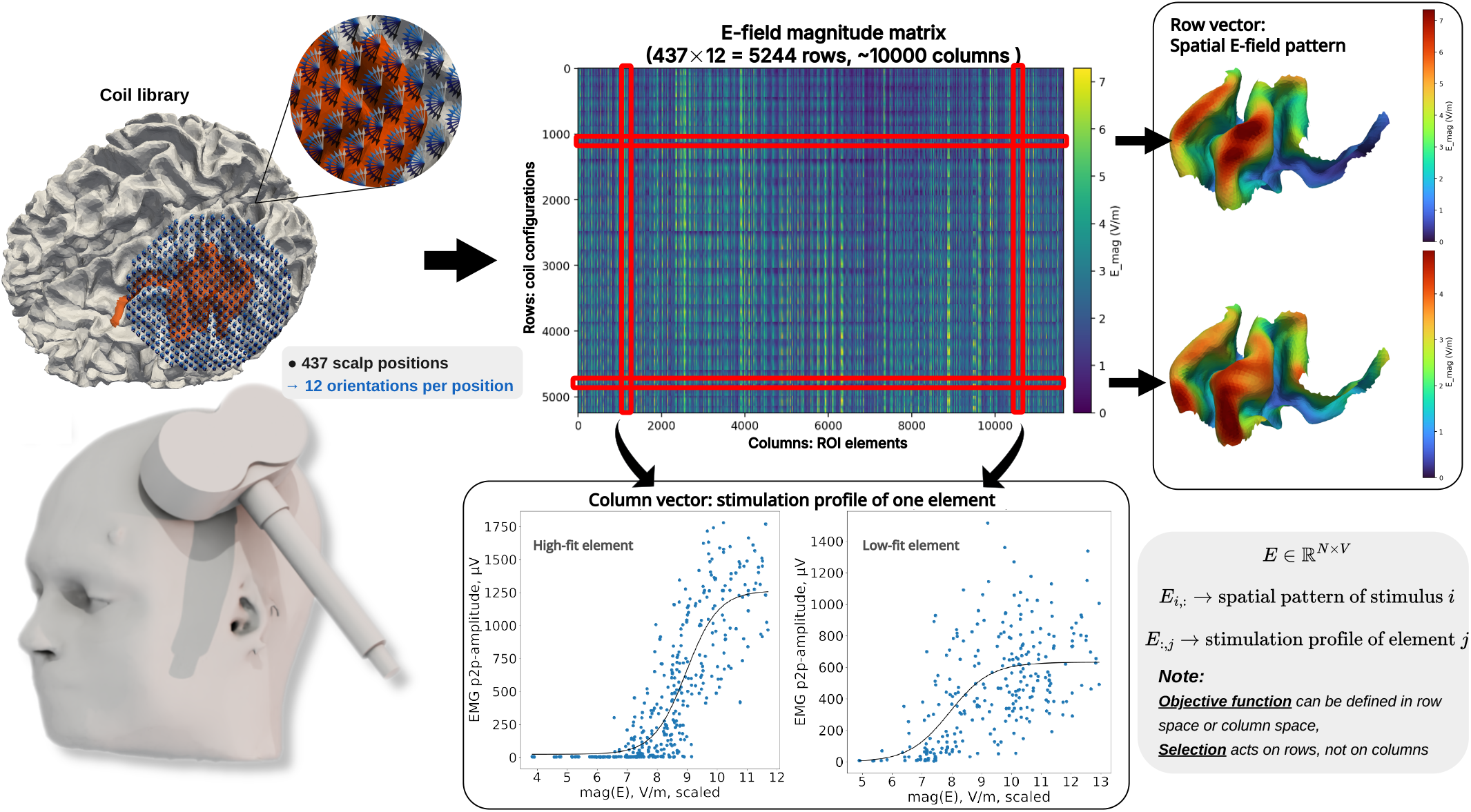
Pattern representation in the unified greedy selection framework for prospective coil-sequence design. The candidate coil library defines an E-field matrix with one row per stimulus and one column per ROI element. Row-wise patterns describe the spatial E-field distribution induced by a single stimulus, whereas column-wise profiles describe how a given cortical element is sampled across stimuli. Although subset criteria may be formulated in either representation, greedy selection always operates on stimuli: each step adds one new row to the design matrix, and column profiles change only indirectly as a consequence of this row addition.

#### E-field extraction

For each candidate coil placement *i*, we extracted the E-field **E**_*i*_(*v*) at each cortical element *v* in the ROI, sampled at the mid-layer of the gray matter to reduce boundary artifacts (Weise et al., 2023). The resulting feature vector **e**_*i*_ ∈ ℝ^*V*^ (where *V* ≈ 10,000 elements per subject) represents the spatial E-field pattern delivered by that coil placement.

#### E-field component selection

All analyses used the E-field magnitude ∥**E**(*v*)∥ at each cortical element. We chose magnitude because it is defined consistently across gyri and sulci, retains amplitude differences, and matches prior E-field-informed motor-mapping work (Jing et al., 2023; Numssen et al., 2021; Weise et al., 2020, 2023); surface-normal and direction-weighted projections were not used.

### 2.3 Unified framework: subset selection with three design choices

An effective prospective coil sequence should avoid redundant placements while favoring E-field patterns that add information and improve element-wise separability across trials. We describe the objective *f* (·) through three design choices (Figure 1): (i) the **pattern representation**, diversity measured either between stimulation-wise patterns **e**_*i*_ ∈ ℝ^*V*^ (rows; do two coil placements induce different fields?) or between element-wise profiles 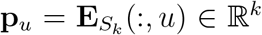 (columns; are two cortical elements distinguishable across the selected stimulations?); (ii) the **pairwise dissimilarity measure**; and (iii) the **aggregator** that converts the pairwise quantities into a set-level score.

We approach prospective coil selection through the lens of **optimal experimental design** (Joshi & Boyd, 2009). Under the general subset objective in Eq. (1), we build sequences with greedy forward selection:

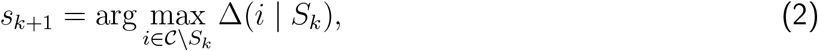

where Δ(*i* | *S*_*k*_) = *f* (*S*_*k*_ ∪ {*i*}) − *f* (*S*_*k*_) is the marginal gain of adding candidate *i* to the current partial set *S*_*k*_ = {*s*_1_, …, *s*_*k*_}. Because exhaustive evaluation over all 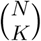 subsets is NP-hard and infeasible at this scale, all methods below use greedy forward selection.

At any step *k*, the selected placements *S*_*k*_ define an E-field “design matrix”

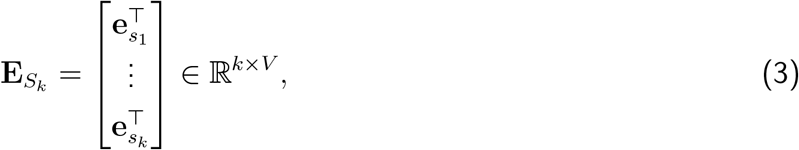

with one row per stimulation and one column per ROI element; each new stimulation appends one row. For the local-separation family, the greedy rule can be written as

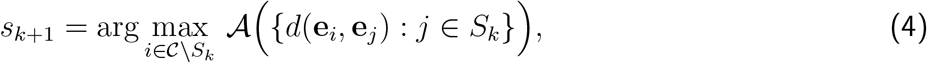

where *d*(·, ·) is a dissimilarity metric and *A*(·) aggregates candidate-to-set dissimilarities (e.g., min for maximin coverage or the mean for average separation). Volume-based objectives (Section 2.3.2) instead aggregate the full similarity structure via a log-determinant. Algorithm 1 summarizes the greedy template for the unified framework.

#### Algorithm 1

Greedy selection template under unified framework

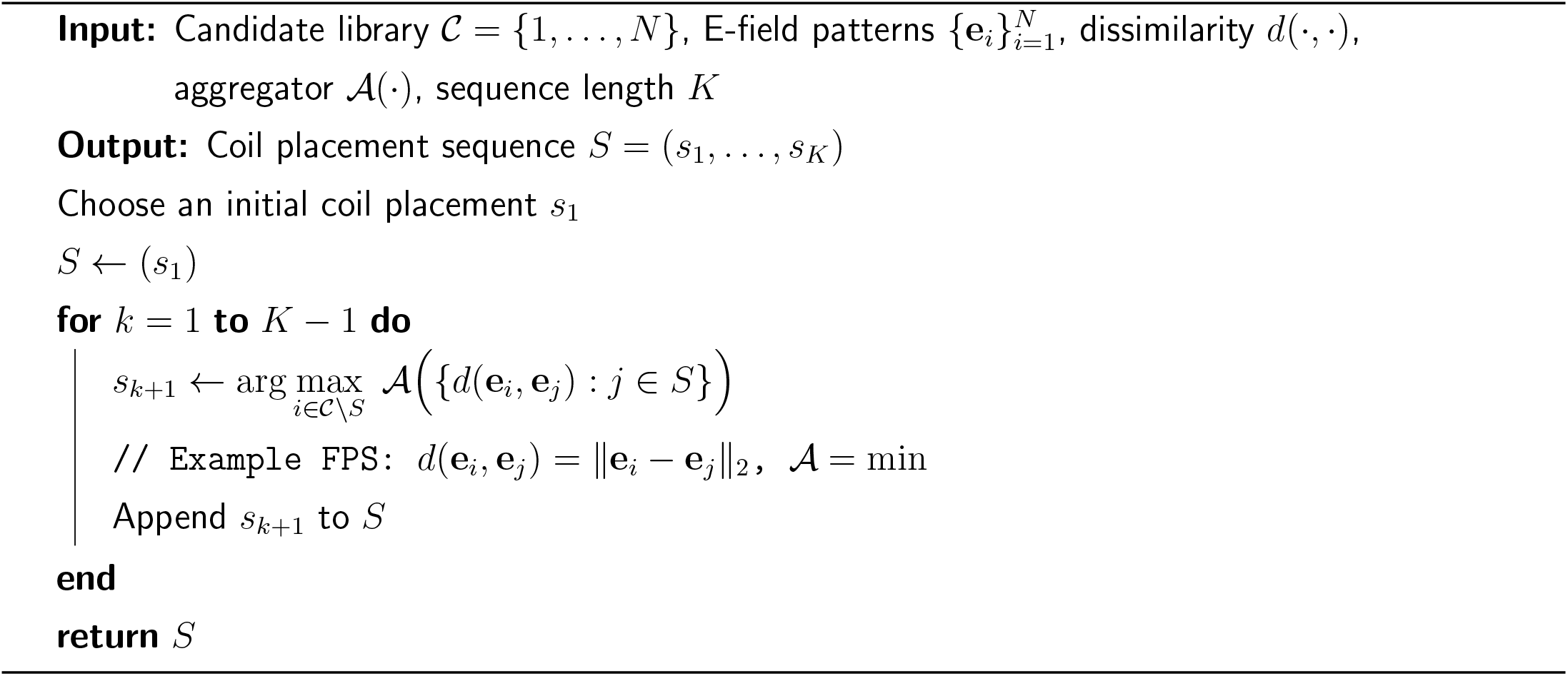

#### 2.3.1 Similarity metrics and local-separation aggregators

In the E-field library, each coil placement corresponds to a high-dimensional pattern vector **e**_*i*_ ∈ ℝ^*V*^ over ROI elements. We quantify redundancy through pairwise dissimilarities between these stimulation-wise patterns.

##### Pairwise dissimilarity metrics

We consider two standard choices. The amplitude-sensitive Euclidean distance,

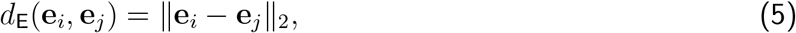

depends on both the angle between patterns and their magnitudes and is unbounded (Figure 2A). In contrast, an amplitude-invariant angular distance based on cosine similarity is

**Figure 2:**
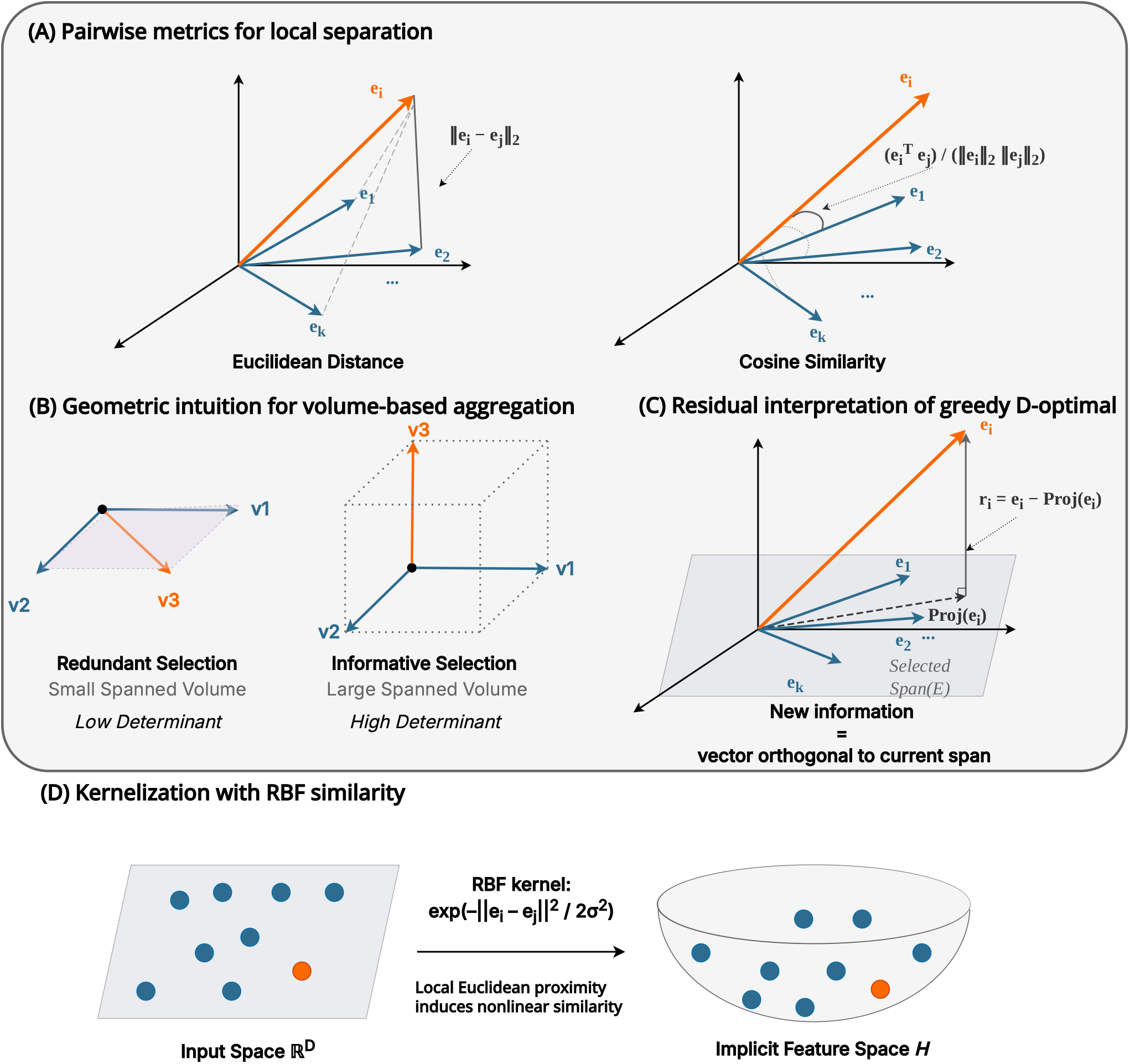
Geometric views of different design choices under the unified subset-selection framework. (A) illustrates alternative choices of pairwise metric for local-separation objectives, using Euclidean distance or cosine similarity between candidate stimulation patterns. (B) and (C) illustrate the volume-based aggregation family: the determinant of the linear Gram matrix summarizes the joint non-redundancy of the selected set, while greedy updates can be understood through orthogonal residual energy relative to the span of previously selected patterns. (D) shows kernelization as a change in similarity representation rather than aggregation.

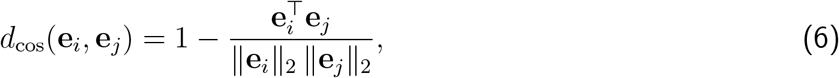

which measures differences in spatial *shape* (Figure 2A) and is invariant to global amplitude scaling. Because we use E-field magnitude (nonnegative entries), cosine similarity is nonnegative in practice, and *d*_cos_ ∈ [0, 1] for our libraries.

Cosine and Euclidean distances become equivalent after *ℓ*_2_ normalization. Let 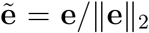 and denote 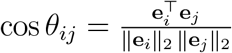 Then

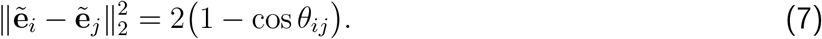

Thus, the key modeling decision is whether distances are computed on raw patterns (amplitude-sensitive) or normalized patterns (amplitude-invariant).

The boundedness of cosine distance has a concrete consequence for the maximin (FPS) aggregator. Because E-field magnitudes are nonnegative and cosine distance satisfies *d*_cos_ ∈ [0, 1], the largest achievable pairwise separation is 1. Euclidean distance is unbounded: two patterns that differ primarily in amplitude can be far apart in *ℓ*_2_ without being distinct in spatial shape. When FPS maximizes the minimum Euclidean distance to the already-selected set, it therefore has an incentive to choose high-amplitude patterns even when those patterns are geometrically similar (in shape) to previously selected ones. Figure 2A illustrates this contrast.

##### Local-separation aggregators

Given *d*(·, ·), local-separation greedy rules instantiate Eq. (4) by aggregating distances from a candidate to the already-selected set. A published work for prospective TMS mapping is **Farthest Point Sampling (FPS)** by Schultheiss et al. (Schultheiss et al., 2025), which uses Euclidean distance with a maximin (min) aggregator:

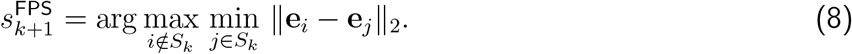

We also consider an average-separation variant that replaces the worst-case criterion with the mean:

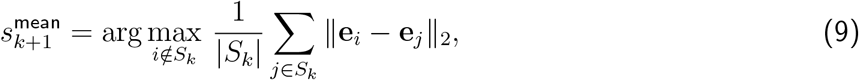

#### 2.3.2 Information-volume aggregation via determinant of the similarity matrix

Local-separation criteria summarize distances from a candidate to the current set via a scalar aggregator (e.g., min or mean). A complementary approach is to penalize redundancy with respect to the entire selected set: a candidate is informative if its stimulation pattern cannot be well approximated by a linear combination of previously selected patterns, as illustrated in Figure 2B. For a selected set *S*, let **E**_*S*_ ∈ ℝ^|*S*|×*V*^ collect the corresponding E-field patterns as rows and define the **Gram matrix**

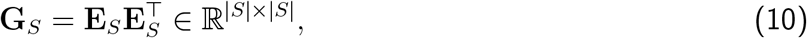

whose entries 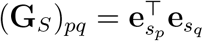 are pattern–pattern inner products. When each pattern is *ℓ*_2_-normalized, (**G**_*S*_)_*pq*_ equals cosine similarity, so **G**_*S*_ can be interpreted directly as a **cosine-similarity matrix** for the selected sequence.

We use the log-determinant of the Gram matrix as a volume-type aggregator. Defining

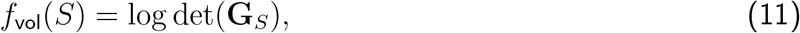

the corresponding D-optimal subset-selection problem is (Joshi & Boyd, 2009; Saito et al., 2021)

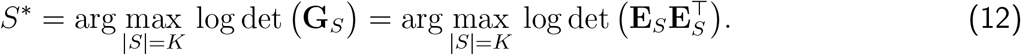

Under the usual linear-model interpretation, maximizing log det(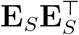) minimizes the determinant of the parameter covariance (Joshi & Boyd, 2009). If the selected patterns are highly redundant, **G**_*S*_ approaches singularity and the spanned volume collapses towards a lower-dimensional subspace. A more balanced spectrum corresponds to a larger parallelepiped volume in the selected-pattern space (Figure 2B). In spectral form,

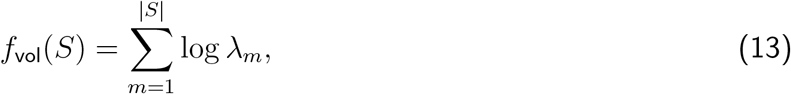

where {*λ*_*m*_} are the eigenvalues of **G**_*S*_.

In greedy forward selection, the marginal gain of adding a candidate can be interpreted as the energy of its component orthogonal to the subspace already spanned by the selected patterns. This connects logdet maximization to pivoted QR / Gram–Schmidt updates and enables efficient implementations (Drmač & Gugercin, 2016; Manohar et al., 2018; Saito et al., 2021). We therefore use stimulation-wise redundancy in the Results as an empirical diagnostic of how this objective behaves in practice.

We use greedy forward selection by choosing the coil that yields the largest increase in log det(**G**_*S*_):

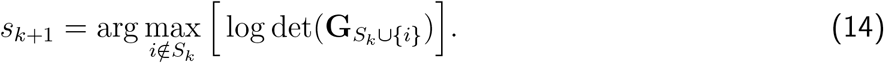

Algorithm 2 provides an equivalent implementation in the sense that maximizing the product of orthogonal residual norms maximizes det(**G**_*S*_). Figure 2C illustrates how the selection criteria for additional coil placements can be interpreted geometrically.

##### Algorithm 2

Greedy linear-kernel D-optimal row selection (max-volume)

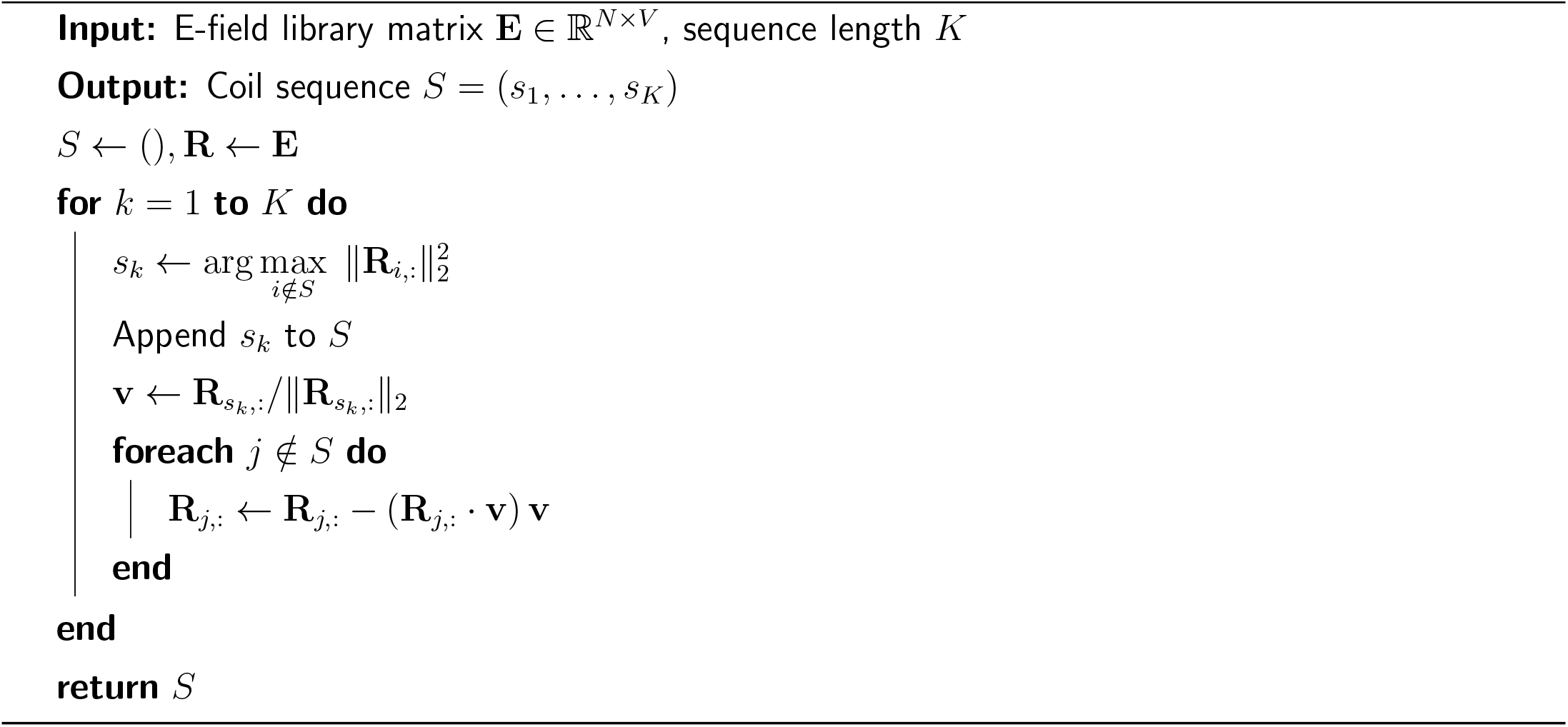

### 2.4 Singular-spectrum structure of the E-field library

We characterize library diversity from the singular spectrum 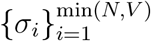 of the E-field library matrix **E**: a steep spectrum indicates stimulation energy concentrated in a few orthogonal directions, while a gradual decay indicates a richer candidate space. For compact reporting and for the subset-level diagnostics below, we summarize the spectrum by its numerical rank, defined as the number of normalized singular values exceeding a tolerance threshold:

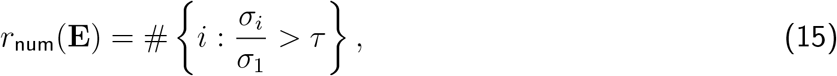

where {*σ*_*i*_} are in descending order. We use the NumPy default scale-aware threshold for single precision, *τ* = max(*N, V* ) · *ε*_*f*32_ with *ε*_*f*32_ ≈ 1.19 × 10^−7^. The spectrum itself remains the more informative object; *r*_num_ is a derived scalar summary.

The candidate library depends on three discretization hyperparameters, spatial step, angular step, and search radius. We use the singular spectrum (with *r*_num_ as a scalar shortcut) as an algorithm-agnostic diagnostic of how each hyperparameter shapes library diversity; Section 3.1 reports this sensitivity analysis before the algorithm comparison.

The steep spectrum motivates non-linear kernelization. Under the linear model, 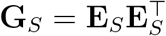 has at most *r*_num_ non-negligible eigenvalues, so after roughly *r*_num_ selections the linear logdet objective in Eq. (12) loses discriminative power: later gains rest on distinctions that the linear similarity model captures only weakly.

To maintain sensitivity to local differences after the dominant spectral subspace is spanned, we keep the same logdet aggregator but define similarity through a nonlinear kernel. We use the radial basis function (RBF) kernel:

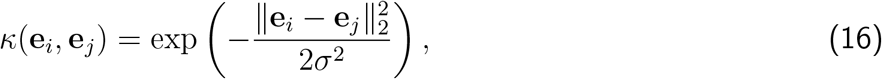

where *σ >* 0 is the bandwidth parameter. The RBF kernel converts Euclidean distance into a bounded similarity as shown in Figure 2D: patterns that are close in Euclidean distance have kernel values near 1, while distant patterns approach 0.

Let **K**_*S*_ ∈ ℝ ^|*S*|×|*S*|^ denote the kernel Gram matrix obtained by replacing each linear inner product in **G**_*S*_ with the RBF similarity from Eq. (16), such that 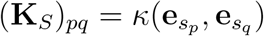 . The kernelized volume objective is therefore

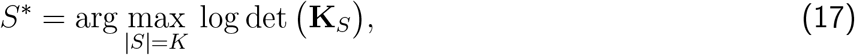

and greedy selection proceeds analogously by maximizing the marginal increase of log det(**K**_*S*_) when adding a candidate coil. We use an incremental Cholesky update to compute the per-candidate marginal gain in *O*(|*S*|^2^) rather than recomputing the full determinant; pseudocode for the kernelized greedy procedure is given in Appendix B.

In the unified framework, this kernelized objective is best viewed as a change of *similarity representation*: we keep the same logdet aggregator but replace the linear inner product with an RBF similarity. The kernel uses Euclidean distance to encode local similarity, while the determinant still measures set-level diversity as a volume.

The bandwidth *σ* controls the scale at which patterns are distinguished. For each subject, we computed Euclidean distances between candidate E-field patterns within that subject’s library and set *σ* to the median of those distances, so the kernel scale adapts to the typical spacing of E-field patterns for that individual library.

### 2.5 Controlled comparisons of pattern representations, metrics, and aggregators

We evaluate a focused set of objectives from this design space (Table 1): random sampling, FPS, FPS-based variants, and two D-optimal variants. All comparisons use the same candidate library and target sequence length *K*.

**Table 1:** Summary of compared methods instantiated within the unified three-choice framework. Methods marked with † are computationally demanding because adding a stimulation changes all column profiles.

| Method | Pattern representation | Metric | Aggregator |
| --- | --- | --- | --- |
| Random | — | — | — |
| Farthest Point Sampling (FPS) | Row-wise | Euclidean ($\ell_2$) | Min |
| corr-FPS | Row-wise | Cosine distance | Min |
| mean-FPS | Row-wise | Euclidean ($\ell_2$) | Mean |
| column-FPS† | Column-wise | Euclidean ($\ell_2$) | Min |
| D-Optimal | — | Linear inner product | Determinant (volume) |
| D-Optimal (kernel) | — | RBF kernel | Determinant (volume) |

**Table 2:**
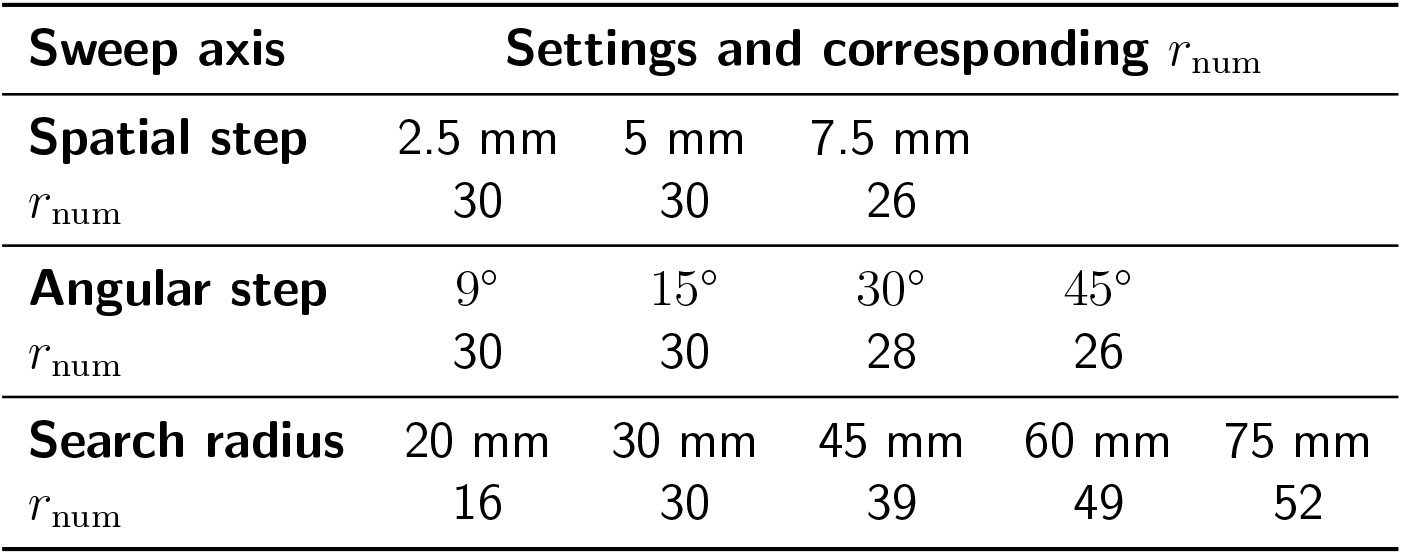
Thresholded numerical-rank summary of the discretization sweep. Numerical rank is reported only as a compact scalar summary of the singular spectrum using Eq. (15); the spectrum shape in Figure 4 is the primary diagnostic.

For the metric, we contrast Euclidean distance on raw patterns with cosine distance. From Eq. (7), the distinction reduces to whether the preprocessing step preserves amplitude information. For the aggregator, we contrast maximin coverage (FPS, corr-FPS; Eq. (8)), mean-distance aggregation (mean-FPS; Eq. (9)), and logdet volume (Eq. (12)): maximin and mean summarize pairwise candidate-to-set distances by an extreme-value or additive statistic, whereas logdet aggregates the full Gram-matrix structure as a volume. The kernelized logdet objective (Eq. (17)) keeps the same volume aggregator but changes the similarity representation.

As a pattern-level control, we also test *column-FPS*, a column-wise decorrelation rule that switches the pattern definition from coil-induced rows to cortical-element columns. With **E**_*S*_(:, *u*) ∈ ℝ^|*S*|^ the stimulation profile of ROI element *u* across the selected set *S*, the next pose maximizes the minimum pairwise Euclidean distance between element-wise profiles:

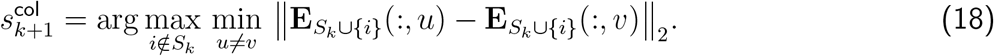

Because adding a stimulation alters all column profiles, this objective is evaluated on a downsampled subset of ROI elements. The control tests whether directly encouraging element-wise separability is more effective than decorrelating stimulation patterns.

### 2.6 Virtual mapping simulations and evaluation metrics

#### 2.6.1 Virtual mapping design

We ran virtual mapping experiments on all 12 individualized head models. For each subject, four ground-truth hotspots were placed within the ROI to represent anatomically distinct contexts, including the gyral crown, the sulcal wall in the fundus of the central sulcus, and peripheral ROI locations. For every (subject, hotspot, method) triple, a prospective coil sequence of length *K* = 100 was generated from the candidate library C (Section 2.2), held fixed, and used to simulate 20 independent realizations of virtual responses, isolating stochastic response variability from differences in sequence design. After the first *k* ∈ {1, …, *K*} pulses, the mapping pipeline yielded a motor map **m**(*k*) over ROI elements and an estimated hotspot Ĵ (*k*) = arg max_*j*_ *m*_*j*_(*k*).

Virtual responses were generated with the dual-variability sigmoidal MEP model adapted from prior TMS mapping work (Goetz et al., 2014; Numssen et al., 2021; Weise et al., 2023), in which the observed MEP at coil configuration *i* depends on the local E-field magnitude at the ground-truth hotspot through a sigmoid with independent input-side and output-side Gaussian noise (Figure 3). Per-subject parameters were held fixed across sequence-design methods and stochastic realizations so that mapping differences reflected the selected E-field sequences rather than changes in the underlying response model; only the sampled noise terms varied across the 20 realizations.

**Figure 3:**
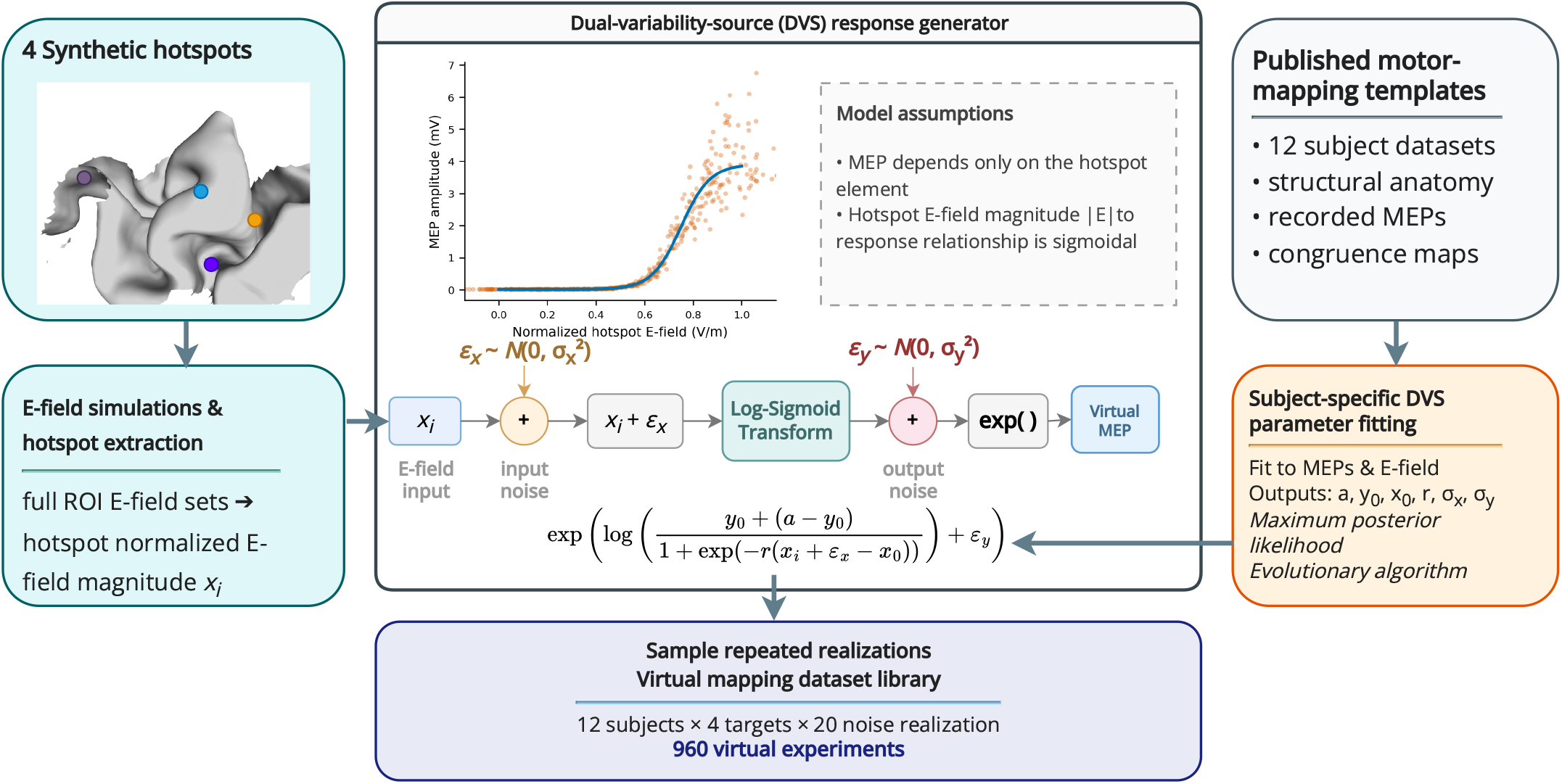
Exemplary virtual MEP datasets generated using the dual variability source model from (Goetz et al., 2014). The inner figure shows the relationship between normalized electric field (in V/m) and MEP amplitude (in mV) for noisy model simulations. The model parameters (e.g. for one subject: *a* = 3.61 ± 2.19 mV, *r* = 16.86 ± 4.96 m/V, *x*_0_ = 0.75 ± 0.12 V/m, *y*_0_ = 8.99 · 10^−3^ ± 13.69 · 10^−3^ mV) were fitted to experimental data from Numssen et al. (2021) by maximizing the likelihood of the posterior distribution. MEP noise was simulated with *σ*_*x*_ = 9.36 ·10^−2^ ± 3.48 ·10^−2^V/m and *σ*_*y*_ = 9.29 ·10^−2^ ± 4.59 ·10^−2^ to mimic realistic conditions and investigate the influence of noise on localization accuracy.

The map value *m*_*j*_(*k*) is the coefficient of determination (*R*^2^) obtained at ROI element *j* by fitting the local E-field–response relationship on the log-transformed simulated MEP responses (Numssen et al., 2021; Weise et al., 2023). The estimated hotspot at each pulse count is thus the cortical element whose local E-field best explains the accumulated observations under the assumed input–output model. The ROI was represented as a triangulated cortical surface with vertex coordinates 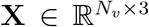 and face matrix **F** ∈ ℕ^*V* ×3^; all analyses were performed at the surface level, with **m**(*k*) ∈ ℝ^*V*^ assigning one scalar per surface element.

#### 2.6.2 Hotspot localization error (geodesic distance)

Hotspot localization accuracy was quantified by the geodesic distance (GD) along the cortical surface between the estimated hotspot element and the corresponding ground-truth element. Because geodesic distances are evaluated on mesh vertices, each surface element was represented by its three constituent vertices. Let *S* = **F**[*j*_true_] denote the set of source vertices belonging to the true hotspot element and *T* = **F**[ĵ] the vertices of the estimated hotspot element. GD was then defined as the mean shortest-path distance from the target vertices to the source set:

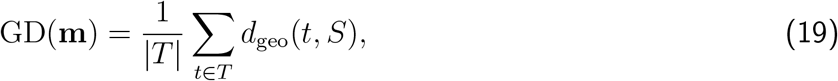

where *d*_geo_(*t, S*) is the shortest geodesic distance from target vertex *t* to the nearest source vertex in *S*. Smaller GD values indicate more accurate hotspot localization.

#### 2.6.3 Map fidelity (normalized RMSD)

Hotspot localization alone does not capture recovery of the full spatial map. We additionally summarized the deviation of the running, max-normalized *R*^2^ map from a single-element binary reference centered at the ground-truth hotspot using the cortical-map NRMSD definition of Numssen et al. (Numssen et al., 2021); the full formula is given in Appendix C. Lower NRMSD indicates closer recovery of the target map.

#### 2.6.4 Convergence summary via AUC

Early mapping efficiency under a clinically relevant pulse budget was summarized by the trapezoidal area under the trajectory over the first 50 pulses, for each outcome *y*(*k*) ∈ {GD(*k*), NRMSD(*k*)}:

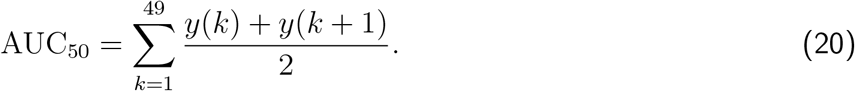

Lower AUC_50_ values indicate more rapid early convergence.

#### 2.6.5 E-field matrix diagnostics

At each pulse count *k*, we evaluated two complementary diagnostics on the selected E-field submatrix 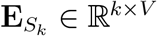. Stimulus-wise redundancy was quantified as the mean absolute Pearson correlation between distinct selected patterns (rows):

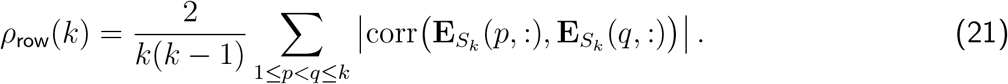

Second, element-wise separability was quantified through the mean absolute Pearson correlation between distinct ROI-element stimulation profiles (columns):

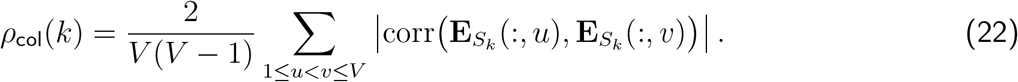

Lower *ρ*_row_ indicates less redundancy among delivered E-field patterns, whereas lower *ρ*_col_ indicates greater separability of cortical elements under the selected sequence. Both diagnostics were analyzed as functions of *k* and summarized using the same hierarchical averaging scheme as the mapping metrics.

#### 2.6.6 Subset-rank recovery diagnostic

As an auxiliary diagnostic of how rapidly each objective recovered the dominant spectral subspace of the full candidate library, we computed the rank gap

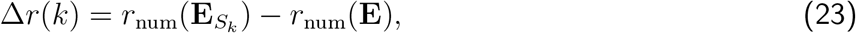

where *r*_num_ is the numerical rank defined in Section 2.4. We evaluated Δ*r* at *k* = 30 and *k* = 100 to compare early and later subspace recovery; interpretive notes on tolerance and conditioning behavior are given in Appendix D.

#### 2.6.7 Low-dimensional embeddings for library geometry

We embedded the row-wise E-field patterns into two and three dimensions using Uniform Manifold Approximation and Projection (UMAP) (McInnes et al., 2018), which builds a high-dimensional neighborhood graph and finds low-dimensional coordinates that preserve local neighborhood structure. The embeddings serve as descriptive geometric summaries, capturing global library organization, method-specific subset coverage in 2D, and intrinsic library geometry under different discretization settings in 3D, and are interpreted jointly with the correlation, rank, and mapping diagnostics rather than in isolation.

## 3 Results

### 3.1 Singular spectrum and intrinsic geometry of the candidate library

The structure of the candidate library constrains how any selection objective can sample it, and prior work has flagged discretization as a robustness concern (Schultheiss et al., 2025). Figure 4 summarizes how the normalized singular spectrum changed when spatial step, angular step, and search radius were varied around the operating point used in the main simulations.

**Figure 4:**
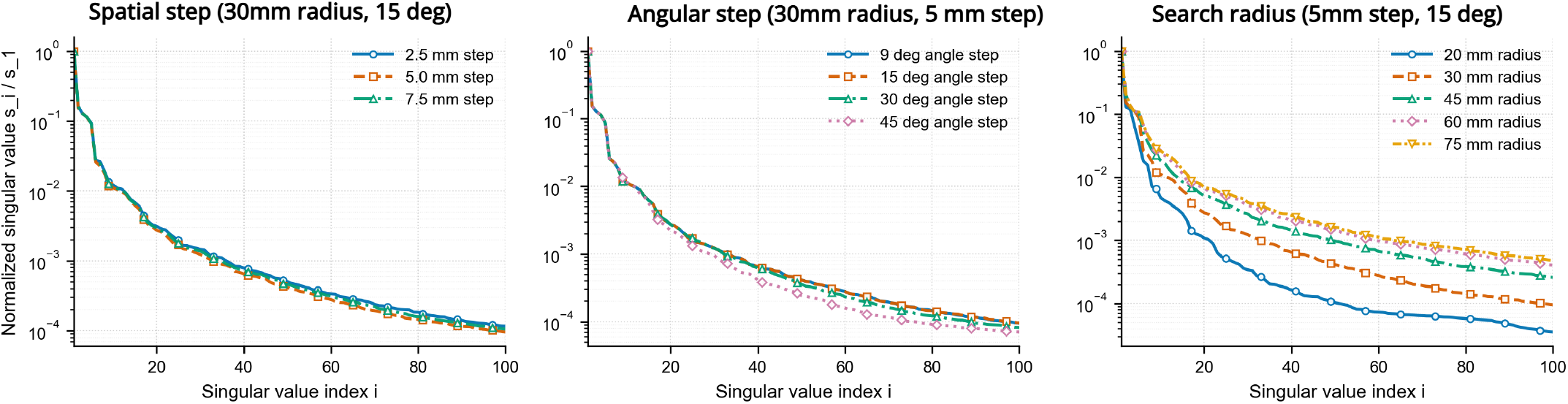
Singular-spectrum sensitivity of the candidate E-field library to discretization hyperparameters. Spectra are normalized by the leading singular value. Spatial-step and angular-step sweeps leave the spectrum shape essentially unchanged across the tested ranges, whereas increasing the search radius from 20 to 75 mm produces a visibly flatter spectrum, with the steepest decay at 20 mm and saturation between 60 and 75 mm.

At the default 30 mm search radius, the spectrum decayed steeply across subjects: *σ*_*i*_*/σ*_1_ dropped below 10^−2^ by *i* ∼10 and below 10^−3^ by *i* ∼40. The spectrum shape was essentially invariant to spatial step across 2.5–7.5 mm and to angular step across 9°–30°; only at 45°did the spectrum visibly steepen, marking the onset of angular coarsening effects. Increasing the search radius from 20 to 75 mm instead flattened the spectrum monotonically, indicating that a larger anatomical search extent contributes additional independent stimulation patterns rather than redundant samples.

As a scalar summary, *r*_num_ changed only modestly across the spatial- and angular-step sweeps but ranged from 16 at 20 mm to 52 at 75 mm across the radius sweep. The SVD diagnostic thus recovers, in a single pass, the same hyperparameter ranking that Schultheiss et al. (Schultheiss et al., 2025, Appendix A.2) obtained by running full FPS-based mapping at every setting.

Whether the added diversity at larger radii translates into better mapping is not automatic. Across our 20/30/45/60/75 mm grid for all algorithms (Appendix F), 20 mm produced the worst GD for every method across both core and periphery hotspots, and worse NRMSD than 30 mm for every method except mean-FPS. The steep 20 mm spectrum anticipates this: the library fell below the diversity floor needed for stable mapping. This supplies a mechanism for the 20 mm-worst / 30 mm-best ordering that Schultheiss et al. reported empirically on a 20/30/40/50 mm grid. The mean-FPS NRMSD inversion is consistent with its peripheral-attraction failure mode (Section 3.3): at 20 mm even “peripheral” placements remain in the strong-field region. At the upper end, spectral gain saturated by 60 mm and brought no convergence improvement; for the magnitude-insensitive corr-FPS objective, performance even degraded with increasing radius, because broader radii admit coil positions producing weak E-fields in the ROI that corr-FPS, ignoring amplitude, cannot down-weight.

The 3D UMAP visualization (Figure 5) gave a complementary geometric view: across subjects, the candidate library formed a smooth manifold rather than a diffuse cloud of independent patterns.

**Figure 5:**
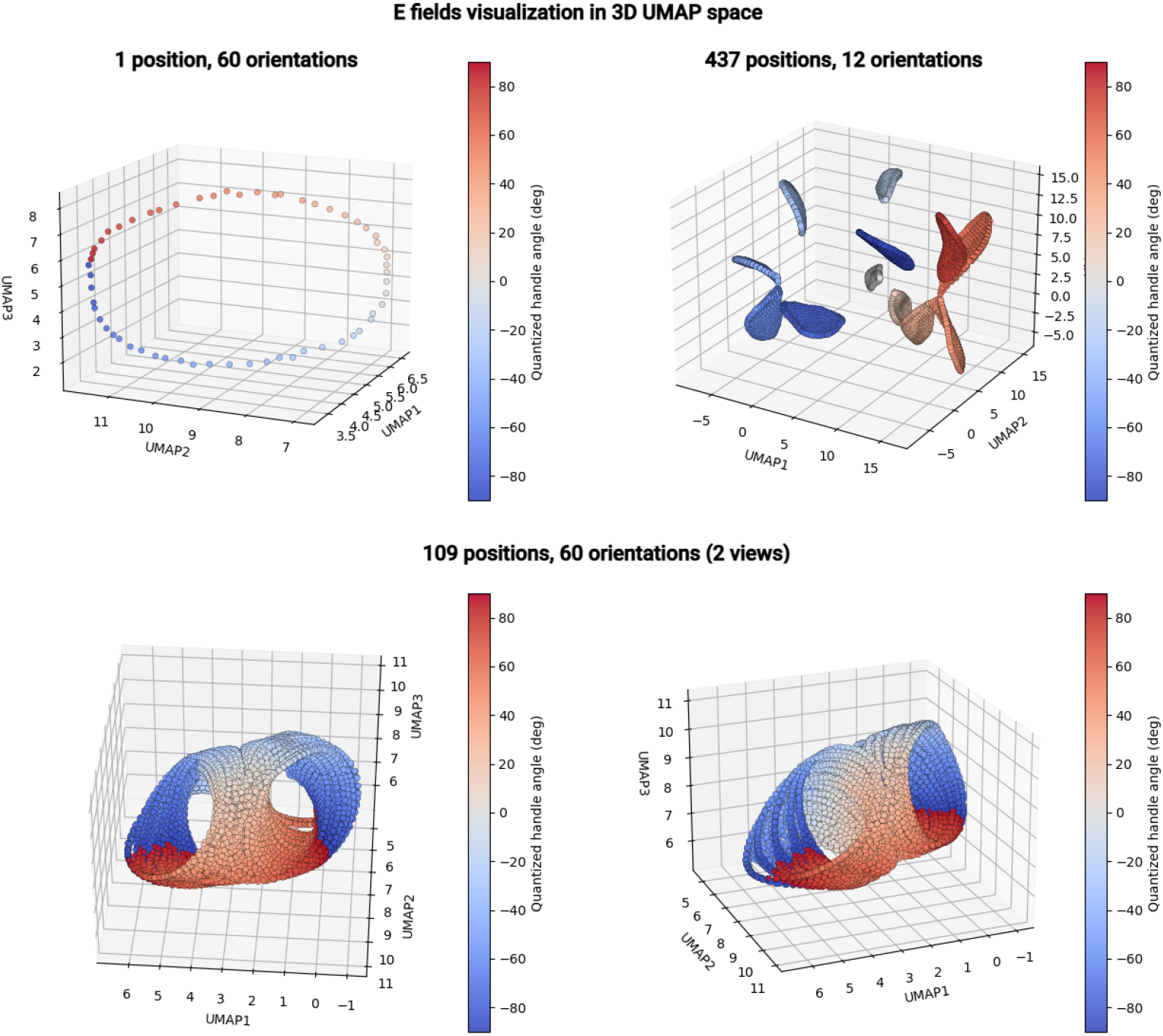
Representative 3D UMAP visualization of the candidate E-field library under different discretization settings. Coil orientation traces a nearly cyclic trajectory; with 12 orientations this appears as orientation-dominated lobes, and denser angular sampling reveals the continuous low-dimensional, non-uniform, partially fused ring-like manifold underlying them. Similar qualitative structure was observed across subjects.

For a single scalp position with 60 orientations, the embedded points formed an approximately closed loop, showing that coil rotation moves the E-field along a nearly cyclic trajectory. With 437 positions and 12 orientations, the embedding separated into roughly 12 leaf-like lobes: orientation set the coarse lobe identity, while position mainly produced within-lobe spread. The angularly denser 109-position, 60-orientation embedding made the underlying continuity more explicit, with distorted and partly fused ring-like ribbons rather than a Cartesian product of position and angle.

Together, the spectrum sweep and 3D UMAP show that the candidate library is a low-dimensional, physics-constrained manifold in which coil rotation contributes more to manifold extent than position change.

### 3.2 Convergence of virtual motor mapping across methods

We next compare selection objectives at the default discretization. Figure 6A shows the full GD and NRMSD trajectories. For all methods, both metrics improved rapidly early in the sequence and then approached a plateau. FPS and both D-optimal variants consistently outperformed random sampling for both hotspot localization and full-map recovery; random sampling converged more slowly and showed broader between-subject variability. The RBF-kernelized D-optimal objective was better during the steep early part of the curves, whereas the linear determinant objective caught up later. The kernelized version also reduced the late NRMSD rebound occasionally seen with the linear design.

**Figure 6:**
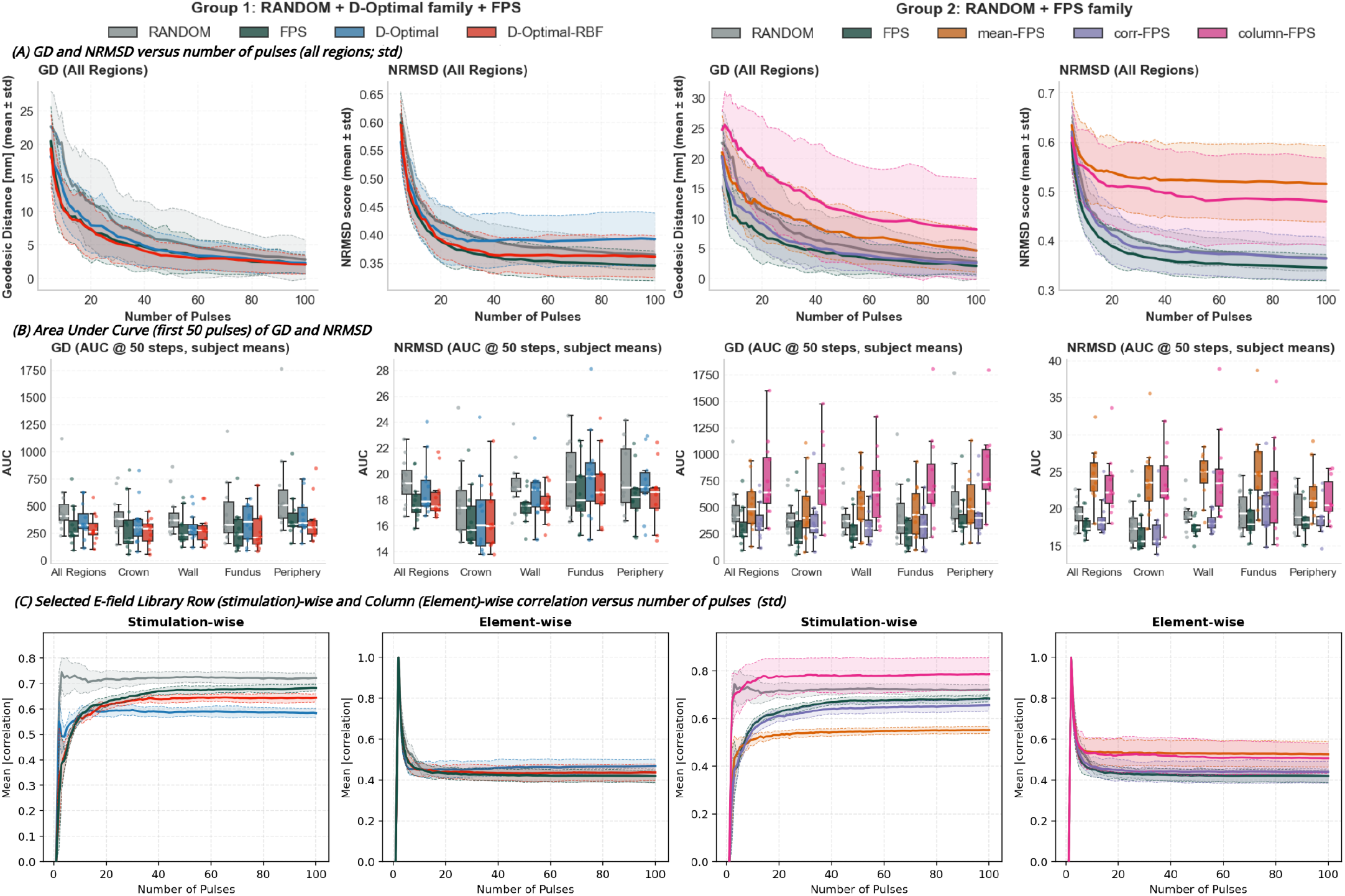
Prospective objectives improve sample-efficient motor mapping, but their benefit depends on the representation and selection criterion. (A) Full convergence trajectories of hotspot localization error (GD) and map fidelity (NRMSD) versus pulse count. Curves denote the mean across subjects after averaging over realizations and hotspot classes; shaded bands indicate standard deviation. (B) Early convergence summarized by the area under the curve (AUC) over the first 50 pulses for GD and NRMSD. The subject-level summaries reproduce the ranking observed in the full trajectories and show that the strongest methods remain favorable across hotspot classes. (C) Matrix diagnostics for the selected E-field submatrices. Better mapping performance is associated with selections that simultaneously limit redundancy among sampled stimulation patterns while maintaining separability across cortical elements, highlighting the importance of balancing these two properties.

Among FPS variants, corr-FPS tracked standard FPS for both GD and NRMSD, indicating that pattern-shape diversity captures most of FPS’s useful non-redundancy, while the amplitude component adds little. Mean-FPS and column-FPS instead underperformed for different reasons: mean-FPS stayed worse than random sampling for GD and fell below column-FPS for NRMSD, while column-FPS was the weakest method for hotspot localization with the broadest between-subject spread.

Subject-level AUC summaries over the first 50 pulses (Figure 6B) preserved this ranking across all regions and within each hotspot class. Both D-optimal variants achieved lower cumulative GD and NRMSD than random sampling, with the kernelized version showing tighter subject-level distributions, particularly for NRMSD. FPS and corr-FPS again yielded the lowest AUCs, while mean-FPS and column-FPS were both worse than random sampling, showing higher AUCs and greater between-subject variability; column-FPS was worse than mean-FPS for GD and comparable for NRMSD.

### 3.3 E-field matrix diagnostics reveal distinct structural trade-offs

We examined the selected E-field submatrix 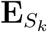 from two complementary viewpoints: stimulation-wise redundancy (mean absolute row-correlation) and element-wise separability (mean absolute column-correlation). Figure 6C shows that methods with similar downstream accuracy could still impose different structural biases on the selected matrix.

Mean-FPS produced the lowest stimulation-wise correlation but the highest element-wise correlation: the pulses became more dissimilar while cortical elements became harder to distinguish across the sequence, matching its GD shortfall relative to random sampling and its NRMSD shortfall below column-FPS. FPS and corr-FPS kept both correlations low and were also the strongest performers on GD and NRMSD. Column-FPS failed on both axes, neither reducing stimulation-wise redundancy nor preserving element-wise separability.

Both D-optimal variants reduced stimulation-wise redundancy relative to random sampling and FPS, consistent with volume maximization penalizing near-collinearity (Section 2.3.2). However, the design acts mainly by expanding the span of selected E-field patterns rather than by directly optimizing element-wise separability, so column correlations were not lower than for random sampling or FPS. The linear variant showed a clearer late increase in element-wise correlation than the kernelized variant, matching its later instability in NRMSD.

As an auxiliary check on these matrix-level findings, we examined how rapidly each method recovered the dominant spectral subspace of the full candidate library through the rank gap Δ*r* (Section 3.1). Linear D-optimal led this diagnostic, FPS and corr-FPS followed closely, and column-FPS recovered the smallest fraction of the dominant subspace; the full method-by-budget summary and Figure 10 are reported in Appendix E. Rank recovery tracked method quality but did not fully predict late map stability: linear D-optimal led on this diagnostic, whereas FPS and kernelized D-optimal were more stable in late map recovery (Figure 6A).

### 3.4 Cross-subject regularities in coil-pose sampling

Different objectives produced systematic regularities in coil-pose selection. Figure 7A summarizes the frequency with which positions and orientations appeared among the first 30 and first 100 selections; because each subject contributed at most one count per position, the position maps reflect cross-subject consensus rather than within-subject reuse.

**Figure 7:**
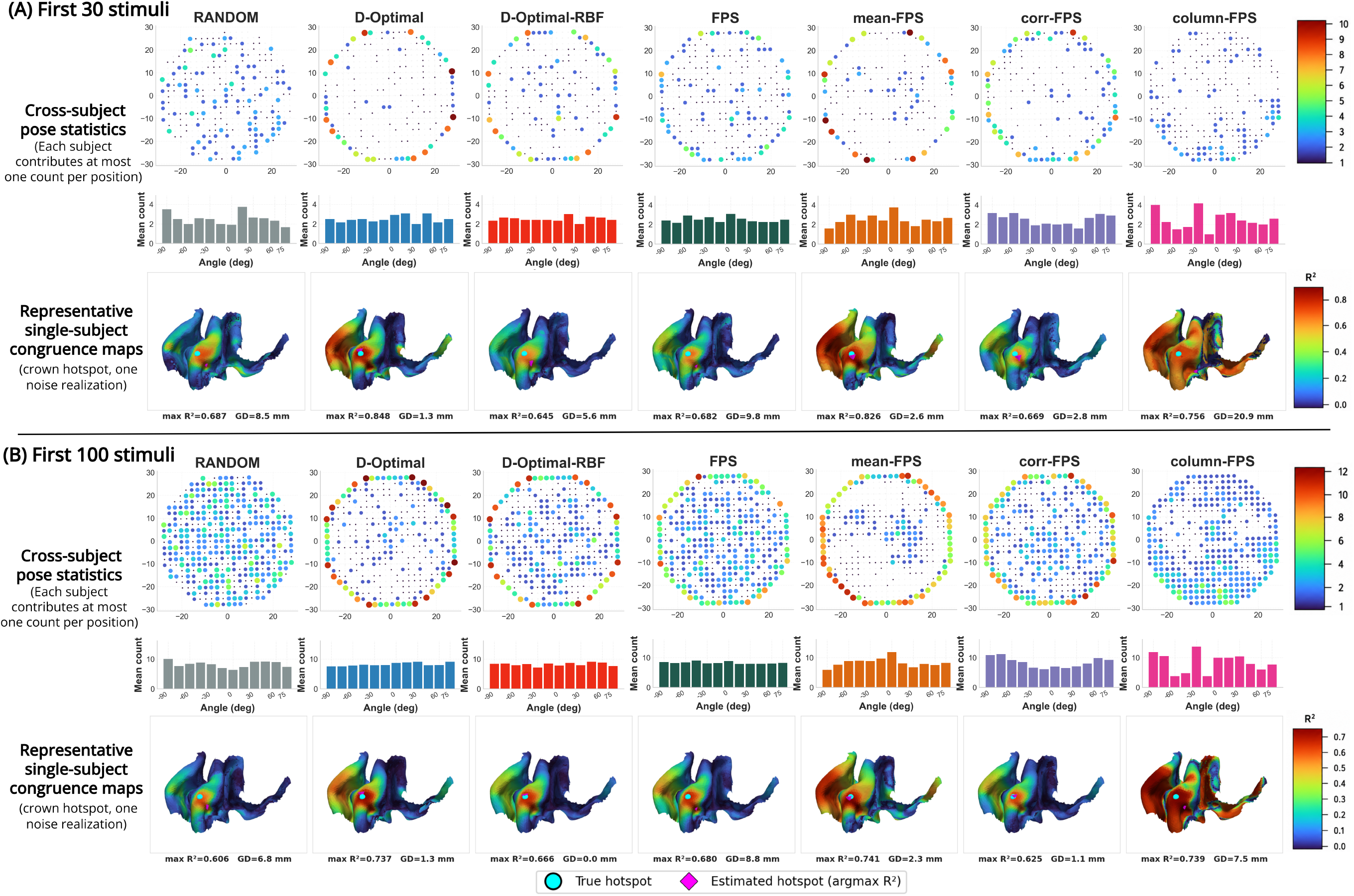
Cross-subject regularities in coil-pose sampling and representative congruence maps after 30 and 100 stimuli. (A) Results after the first 30 selected stimuli. For each method, the top row shows cross-subject position-frequency maps of selected coil poses, the middle row shows the corresponding orientation histogram, and the bottom row shows a representative single-subject congruence map for a virtual subject with a crown hotspot under one noise realization. (B) Results after the first 100 selected stimuli. Congruence maps are shown as element-wise *R*^2^ maps reconstructed from the corresponding selected stimuli. The maximum *R*^2^ and geodesic distance (GD) between the true and estimated hotspots are reported below each map.

Stronger row-wise methods converged on a boundary-first spatial strategy. Early in the sequence, FPS and both D-optimal variants sampled peripheral positions more often, consistent with rapid expansion over the target region: linear D-optimal showed the strongest boundary emphasis, FPS was more balanced, and the kernelized variant fell between them. Corr-FPS resembled standard FPS, mean-FPS stayed boundary-weighted for longer, and column-FPS instead selected interior positions with denser local reuse, suggesting local filling rather than global expansion. The orientation histograms followed the same pattern: better-performing methods used all 12 tangential angles evenly, whereas column-FPS was again less balanced. These differences persisted through 100 selections, with linear D-optimal additionally expanding into the interior while retaining boundary coverage. The illustrative congruence maps in Figure 7 show that broader boundary coverage coincided with earlier focalization and more accurate hotspot localization, whereas denser interior filling could still yield diffuse or displaced maxima at the same sequence length.

Because prospective sequences are computed offline, they can be rearranged for execution by grouping orientations at each scalp position. On neuronavigated TMS-cobot platforms, orientation changes at a fixed contact point require smaller robotic adjustments than transitions between positions (Agboada et al., 2023), so we used the number of unique coil positions as a pragmatic surrogate for movement cost.

Figure 8 shows clear differences in unique-position usage over the first 100 stimulations. Random sampling had the highest positional diversity (87.33 ± 4.11), followed closely by FPS (84.25 ± 3.03) and corr-FPS (83.08 ± 2.72). Linear D-optimal (62.08 ± 6.49) and mean-FPS (63.83 ± 7.00) reused positions much more aggressively, whereas the RBF-kernelized D-optimal method occupied an intermediate regime (69.58 ± 2.22). Column-FPS showed moderate positional diversity (75.33 ± 11.08) but also the largest between-subject variability.

**Figure 8:**
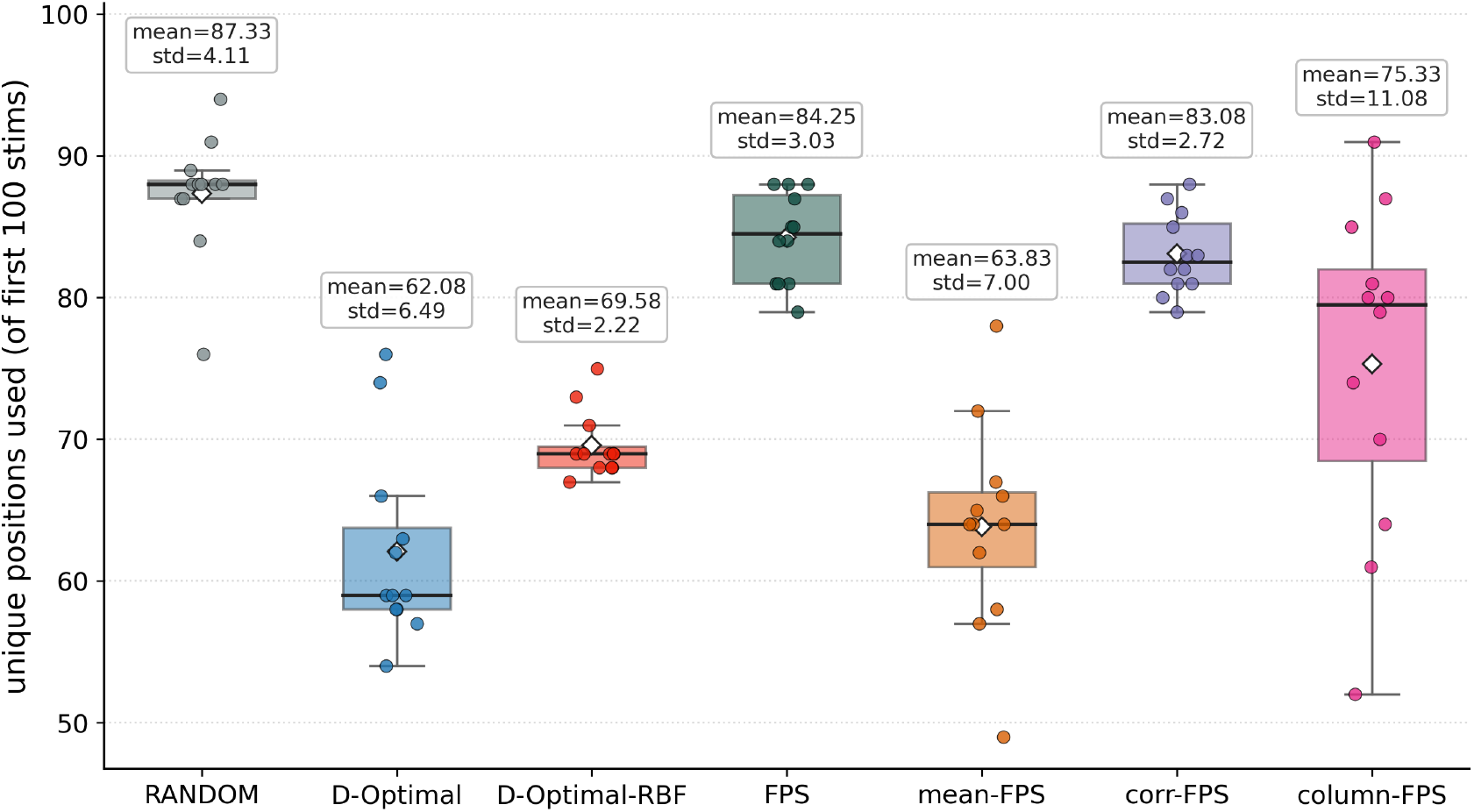
Unique coil positions among the first 100 stimulations after grouping across orientations. Lower values indicate greater positional reuse and therefore lower expected coil repositioning cost.

Among the high-performing methods, the RBF-kernelized D-optimal rule matched FPS in mapping quality (Section 3.2) while using about 15 fewer unique positions. Thus, determinant-based selection reduced repositioning demand without sacrificing mapping quality.

### 3.5 Two-dimensional UMAP shows differential manifold coverage

We then used 2D UMAP to examine whether the selection rules covered the same regions of the candidate E-field manifold. Figure 9 visualizes how the first 100 selections from each method occupy the embedded candidate library. The candidate library was not a uniform cloud; it separated into about 12 lobed regions with lower-density gaps between them. This coarse organization matched the 12-orientation discretization used in the simulation libraries, suggesting that orientation determined the coarse location of a pattern in the embedding, whereas position mainly produced within-lobe variation. FPS, corr-FPS, and both determinant-based methods distributed their selections across most of these regions, consistent with broader manifold coverage. Mean-FPS also covered multiple lobes, but less evenly. Column-FPS showed the clearest collapse, concentrating within only a few local regions and leaving much of the embedded library under-sampled.

**Figure 9:**
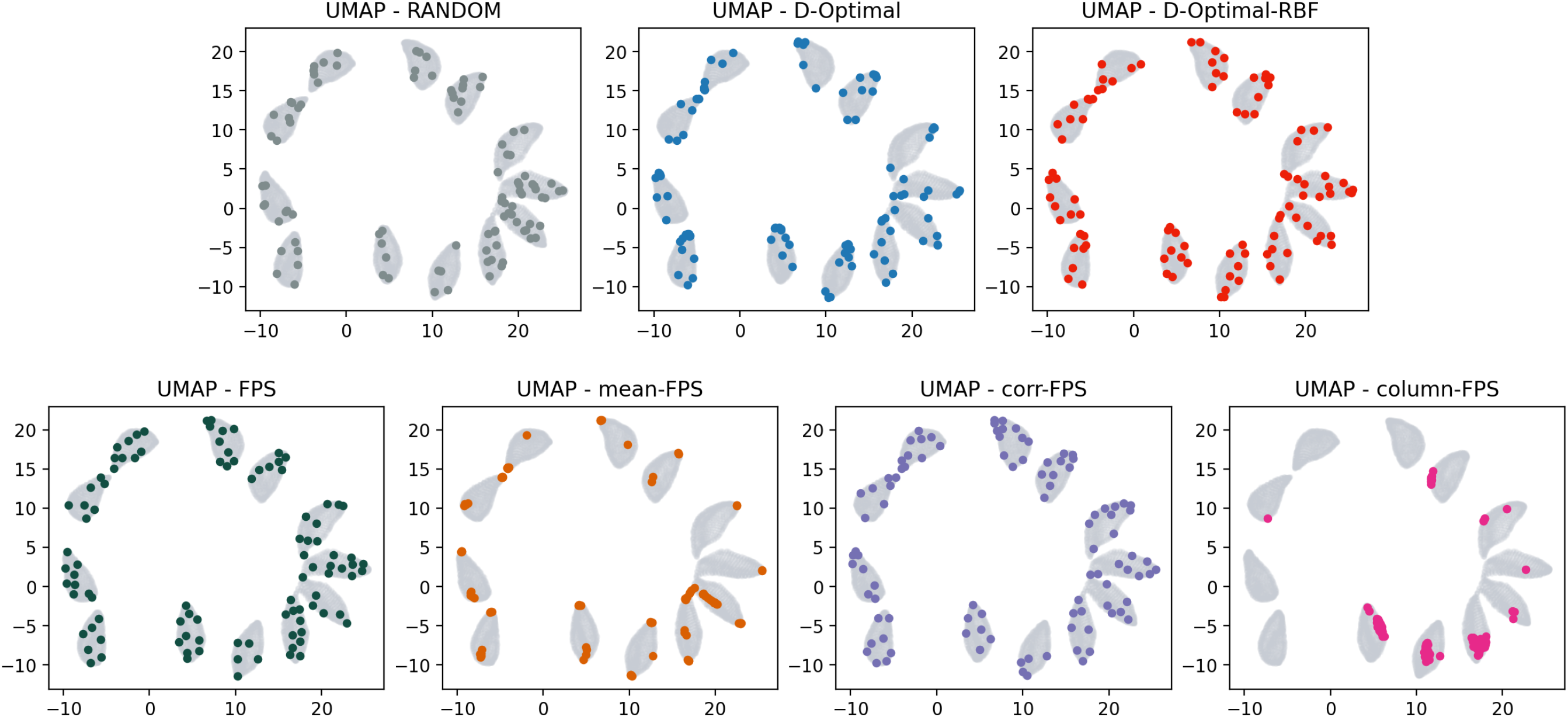
Method-specific coverage of the candidate E-field manifold in 2D UMAP space for the first 100 steps. The major lobes correspond primarily to the 12 simulated orientations, with positional variation producing smaller within-lobe spread. High-performing methods distribute selections across these lobes, whereas column-FPS concentrates on a restricted subset of the library.

Viewed together with the spectrum characterization, subset-rank recovery, correlation diagnostics, and mapping trajectories, these visualizations support the same interpretation: better-performing objectives distributed their selections broadly over the feasible manifold, whereas weaker objectives remained confined to narrower local neighborhoods.

## 4 Discussion

### 4.1 Main findings and interpretation

Prospective coil-sequence design improved the efficiency of E-field-informed motor mapping, but not every diversity objective improved mapping convergence. FPS, corr-FPS, and the determinant-based objectives consistently outperformed random sampling, whereas mean-FPS and column-FPS did not. Across convergence metrics, matrix diagnostics, pose statistics, and UMAP coverage, the strongest methods combined low stimulation-wise redundancy with preserved cortical separability.

### 4.2 Singular-spectrum structure and its hyperparameter sensitivity

The steep spectral decay and the cyclic per-position orientation loop revealed by UMAP both reflect a physically constrained, low-dimensional stimulation space. Selection therefore operates on a curved manifold whose geometrical distribution is constrained by Maxwell’s equations and scalp anatomy before any optimization criterion is applied.

The singular spectrum is not only descriptive of the candidate library but also predictive of the radii at which mapping itself degrades. At the lower end, the collapsed 20 mm spectrum coincided with method-agnostic GD degradation, indicating that the library had too little E-field diversity for reliable hotspot localization. At the upper end, the spectral gain saturated between 60 and 75 mm and did not produce a commensurate mapping gain; instead, broader radii admitted coil positions producing weak E-fields in the ROI that particularly harmed magnitude-insensitive selection such as corr-FPS. Thus, the spectrum predicts both sides of the useful operating range: radii below the diversity floor fail by under-sampling the attainable field space, whereas radii beyond the saturation regime add candidates that are geometrically distinct but not necessarily useful for evoking informative responses.

The same spectral structure explains the early strength and later instability of the linear determinant objective. Maximizing linear information volume efficiently suppresses redundancy and spans the dominant subspace, but once that span is filled the linear criterion has little leverage left for ranking candidates that matter for stable map recovery, consistent with its late rise in element-wise correlation and NRMSD rebound. The RBF kernel does not create new physical degrees of freedom; it reshapes similarity so that nearly collinear patterns can still differ under a localized measure, keeping the kernel objective discriminative after the linear criterion has saturated. The subset-rank diagnostic corroborates this picture: linear D-optimal closed the rank gap to the full library most quickly (especially within the first 30 selections), yet kernelized D-optimal and FPS remained more stable in late map recovery. Fast subspace filling thus helped but was not sufficient, because the candidate library forms a curved manifold rather than a purely linear space.

### 4.3 Why mean-FPS and column-FPS underperformed

Mean-FPS and column-FPS underperformed for different reasons. By construction, mean-FPS reduced row-wise correlation among selected E-field patterns most aggressively. Yet this did not improve mapping; instead, it produced the worst element-wise correlation profile, indicating that the accumulated sequence made cortical elements less distinguishable from one another. Stimulation-wise decorrelation alone is therefore not a sufficient design target in this setting.

Column-FPS fails for a more structural reason. In the row-wise setting, each new stimulation adds one row while leaving previously computed row–row relationships intact; in the column-wise setting, every new stimulation alters all element-wise response profiles simultaneously, so the greedy objective is chasing a moving target. In our data, this instability produced neither useful element-wise decorrelation nor reduced stimulation-wise redundancy. Column-FPS instead selected more interior positions, used the 12-angle set less evenly, and concentrated on a narrower subset of the embedded library.

### 4.4 Practical implications for stimulation planning

The cross-subject pose statistics suggest a practical heuristic for manual mapping protocols. The stronger methods started with broader peripheral or boundary sampling of the available coil placements and only later filled the interior. In these simulations, peripheral placements produced more dissimilar E-field patterns and therefore expanded spatial support more quickly, whereas dense interior sampling was more likely to revisit nearby placements that added little new information. In settings without robotic positioning, early spatial spread may therefore be preferable to dense interior sampling, although this guidance still requires in vivo validation.

The discretization analysis provides a complementary planning heuristic. For this ROI and coil model, spatial steps up to 7.5 mm and angular steps up to 30° remained in the spectrum-stable range, making them reasonable starting points before end-to-end validation. Search radius should match the anatomical extent of the target region, because it materially reshapes the spectrum. In our data, 30 mm occupied a useful middle ground: it cleared the diversity floor that 20 mm failed to clear, while avoiding the diminishing-returns and weak-field regime from 45 mm upward. The SVD spectrum therefore provides a cheap a-priori indicator that the chosen radius spans the relevant E-field diversity without admitting many coil positions producing weak E-fields in the ROI. Within this stable range, coarser settings can substantially shrink library size and the associated simulation cost without appreciably changing the attainable stimulation space.

Position reuse adds a separate practical dimension. Robotic TMS systems deliver accurate placement, but execution cost still scales with how often the system must reposition across scalp sites (Goetz et al., 2019; Lancaster et al., 2004). Offline sequences can be rearranged by grouping orientations at each position, so the number of unique positions approximates the number of full arm reconfigurations. The RBF-kernelized D-optimal method used ∼15 fewer unique positions than FPS within the first 100 stimulations while matching its mapping quality, suggesting a movement-cost reduction without accuracy loss, which is of special interest for robotic TMS applications where arm movement is time-costly.

A method-level screen complements the SVD-based library screen above: the rank-gap diagnostic Δ*r*, computed directly from the candidate library and a candidate sequence (Appendix E), can rule out obviously redundant objectives before any pulse-level simulation, although it does not replace task-level validation.

### 4.5 Limitations and future directions

The present study has several limitations. All comparisons were performed in a virtual mapping framework with subject-specific head models and an established E-field-informed mapping pipeline (Numssen et al., 2021; Weise et al., 2020, 2023). Stochastic MEP variability was modeled explicitly and held constant across methods, isolating the design phenomena, but navigation error, coil-placement imprecision, state-dependent corticospinal excitability, and non-stationary MEP fluctuations were not captured. The position-reuse advantage of kernel D-optimal is similarly a simulated movement-cost estimate rather than a measured one. Within-subject head-to-head comparisons of FPS, the D-optimal variants, and random sampling on a neuronavigated TMS-cobot platform (Agboada et al., 2023; Krieg et al., 2014; Rossini et al., 2015), under matched pulse budgets and real-time MEP acquisition, would convert these estimates into measured session-time reduction and supply the missing in vivo evidence.

The virtual responses assumed a single cortical hotspot, whereas empirical motor maps show spatially extended and overlapping muscle representations (Classen et al., 1998; Jin et al., 2023; Wilson et al., 1993). The manifold-coverage framing developed here is response-model agnostic, so coupling it with forward models that represent extended representations would test whether the diversity-plus-separability rule survives a more physiologically realistic readout, and could also clarify how much of the late NRMSD instability seen with linear D-optimal is intrinsic versus an artifact of the single-hotspot assumption.

Results were obtained with a fixed pulse budget, and greedy forward selection, so the objectives are heuristic rather than globally optimal; ROI definition also enters indirectly, because the ROI determines the candidate library and thus the singular spectrum that any method must cover. A future direction is to extend the present offline design to an online closed-loop setting, where the accumulated MEP evidence is incorporated into the selection process and used to update the priority of the remaining candidate coil configurations during the session, connecting prospective sequence design to the broader move toward closed-loop, brain-state-dependent TMS (Bergmann, 2018).

### 4.6 Conclusion

Prospective coil-sequence design is best framed not as a generic search for decorrelation but as limited coverage of a low-dimensional E-field manifold whose geometrical distribution is constrained by Maxwell’s equations and head anatomy. Within this framing, FPS (Schultheiss et al., 2025) and the determinant-based objectives succeed because they balance two complementary requirements, spreading stimulation patterns apart and at the same time keeping cortical elements distinguishable. In geometric terms, they embody an implicit manifold-coverage rule: by pushing successive stimulations apart in E-field space, they spread samples over the attainable manifold. Our contribution is to make this view explicit and to show why it works. We also provide a singular-spectrum view, indicating which discretization parameters on coil placements are worth probing at all, sidestepping the otherwise expensive sweep of running the full mapping pipeline at every setting. Together with the position-reuse advantage of the kernelized determinant objective, these results give prospective TMS mapping a principled, library-level handle that should make in vivo evaluation cheaper to design, easier to compare across labs, and faster to extend beyond the motor system.

## Data and Code Availability

The algorithm implementations of the prospective coil-sequence selection methods evaluated in this work are available as part of the open-source `pynibs` toolbox at https://gitlab.gwdg.de/tms-localization/pynibs/-/tree/dev/pynibs/optimization. The subject-specific finite-element head models were constructed with SimNIBS 4.5 (https://simnibs.github.io) from anonymized T1- and T2-weighted MRI scans of 12 individuals; these previously acquired experimental data were part of an earlier study (Numssen et al., 2021).

## Author Contributions

Conceptualization: RQ, ON, KW, and TRK; Methodology: RQ, ON, BK, KW, and TRK; Software: RQ, ON, BK, and KW; Formal analysis: RQ; Investigation: RQ, ON, and KW; Data curation: ON and KW; Resources: ON, BK, KW, and TRK; Visualization: RQ; Supervision: KW and TRK; Project administration: TRK; Funding acquisition: KW and TRK; Writing – original draft: RQ; Writing – review and editing: all authors. All authors discussed the results and approved the final manuscript.

## Acknowledgements

The authors thank the developers and maintainers of SimNIBS and `pynibs`, which provided the computational infrastructure for subject-specific electric-field modeling and analysis. The authors also acknowledge the participants whose anonymized MRI data were used to construct the finite-element head models analyzed in this study. The academic English of parts of this manuscript was improved with the assistance of ChatGPT 5.5. The study design, analyses, interpretation of results, and scientific conclusions were entirely the work of the authors.

## A Tissue Conductivity Values

The finite-element head models used tissue conductivity values from the SimNIBS configuration. The assigned conductivities were 0.126 S/m for white matter, 0.275 S/m for gray matter, 1.654 S/m for cerebrospinal fluid, 0.008 S/m for compact bone, 0.025 S/m for spongy bone, 0.465 S/m for skin, 0.600 S/m for blood, 0.160 S/m for muscle, and 0.500 S/m for eyeballs.

## B RBF-kernelized D-optimal greedy selection

Algorithm 3 gives pseudocode for the kernelized D-optimal greedy selection rule defined in Eq. (17). Each iteration extends a running Cholesky factor of the kernel Gram matrix by one row/column so that the marginal-gain proxy 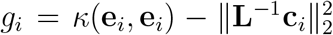 can be evaluated for every remaining candidate at cost *O*(|*S*|^2^), rather than recomputing log det(**K**_*S*_) from scratch at each step.

### Algorithm 3

RBF kernelized D-optimal greedy selection

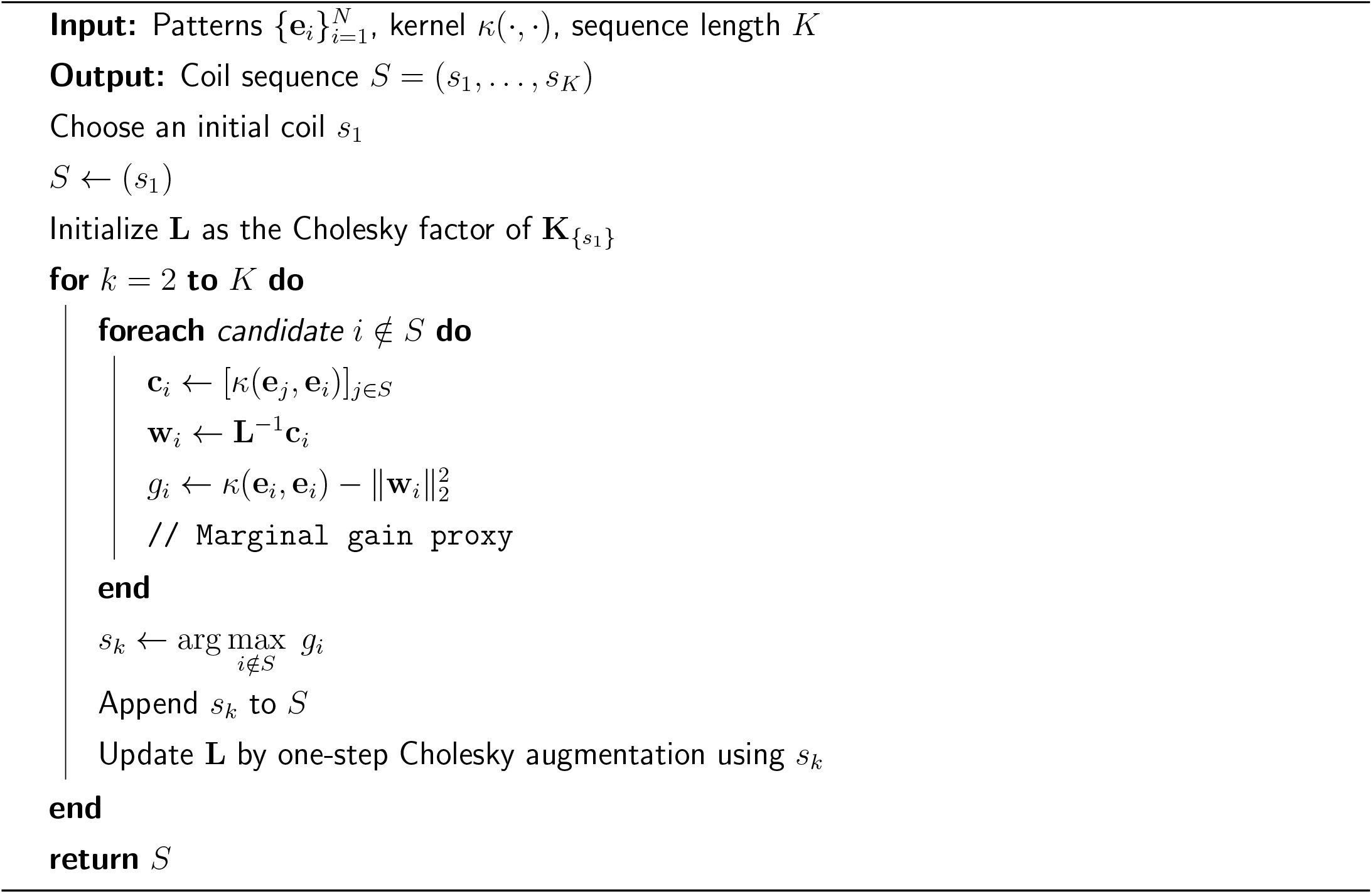

## C Normalized RMSD definition for cortical maps

Hotspot localization alone does not capture recovery of the full spatial map; we therefore additionally compared the running *R*^2^ map to a single-element (“Dirac”) reference 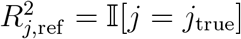, which places all reference mass at the ground-truth element. For each running map after *n* stimuli, the element-wise *R*^2^ scores were first normalized by their maximum value,

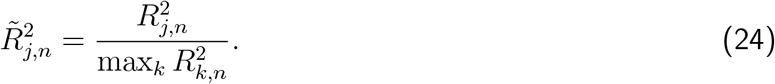

Map deviation was then summarized by the normalized root-mean-square deviation (NRMSD), following the cortical-map NRMSD definition (Numssen et al., 2021):

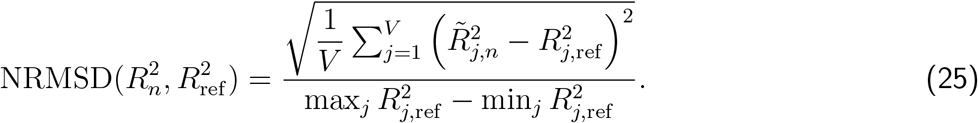

In the present virtual experiments, the reference map is binary with one nonzero element, so 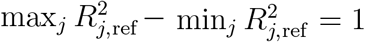. Lower NRMSD indicates closer recovery of the target map.

## D Subset-rank recovery diagnostic: tolerance and interpretation

The rank gap 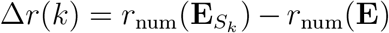 defined in the main text uses the numerical-rank summary *r*_num_ from Section 2.4, which counts the normalized singular values exceeding *τ* = max(*N, V* ) · *ε*_*f*32_ with *ε*_*f*32_ ≈ 1.19 × 10^−7^. Three interpretive notes apply when comparing Δ*r* across methods. Negative values indicate that the selected subset does not yet span the dominant singular directions of the full library, i.e. incomplete subspace recovery. A value of zero indicates that the subset matches the full-library rank summary under the chosen tolerance. Small positive values do not correspond to new physical degrees of freedom, because rank 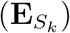 ≤ rank(**E**) in exact arithmetic; instead, they reflect cleaner numerical coverage of the dominant directions when the selected submatrix is better conditioned than the full library under the same tolerance.

## E Subset-rank recovery across methods

We use the rank gap Δ*r* (defined in Section 3.1 and Appendix D) as a scalar proxy for how much of the dominant spectrum a selected subset has captured. Figure 10 summarizes this diagnostic at 30 and 100 selections. Linear D-optimal led: by 30 selections, its rank gap was often at or above zero, indicating the fastest recovery of the major spectral directions. The kernelized variant was slightly less aggressive (gap near zero at 30, positive but smaller than linear D-optimal at 100). FPS and corr-FPS formed a second tier, mildly negative at 30 and near or above zero by 100, consistent with efficient subspace coverage despite an objective that is not explicitly rank-maximizing.

**Figure 10:**
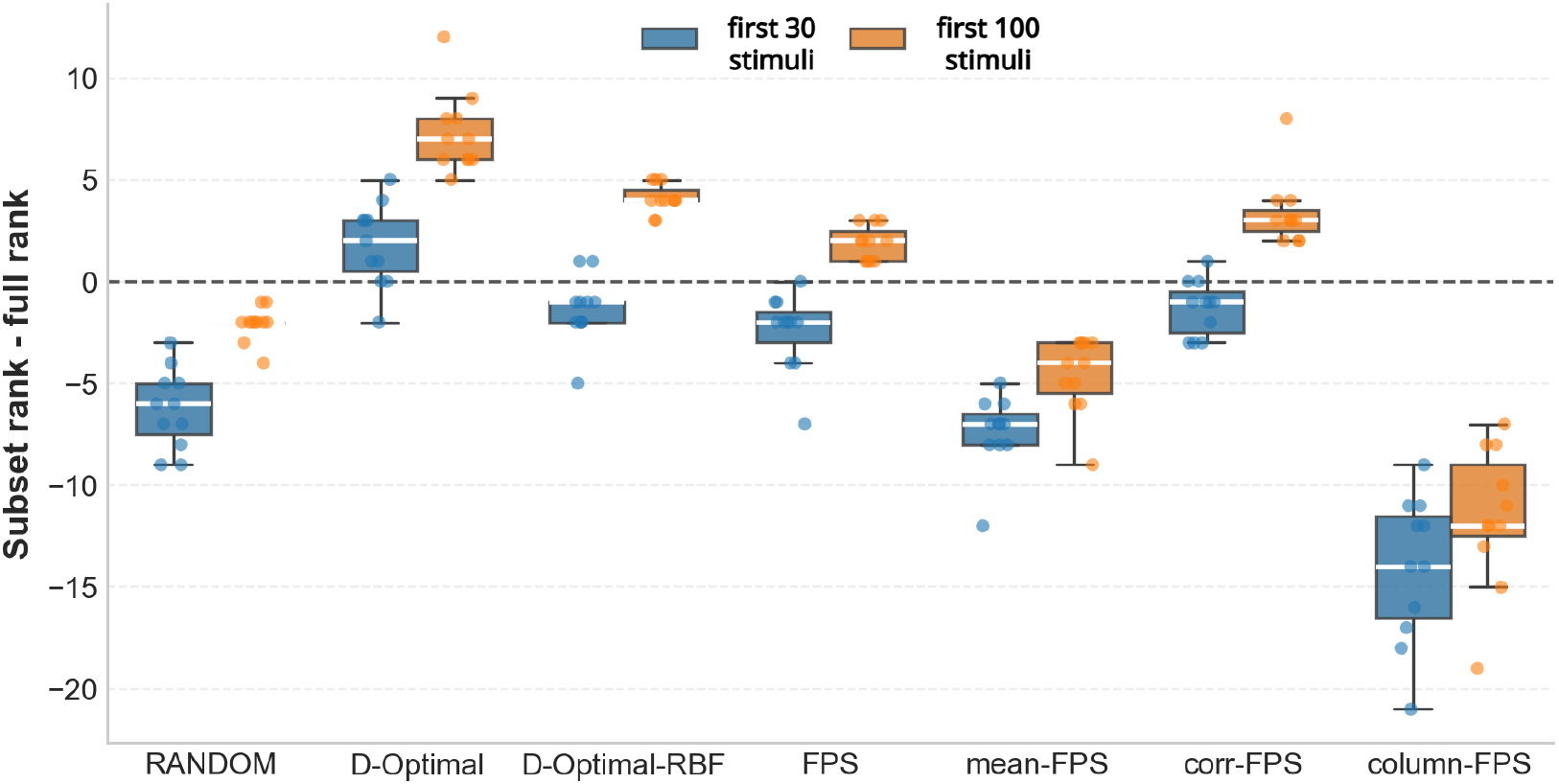
Rank-gap summaries for method-specific first-30 and first-100 stimuli. Metric: 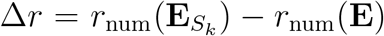 Linear D-optimal most rapidly recovers the thresholded rank summary of the full library, FPS and corr-FPS follow closely, whereas Random, mean-FPS, and especially column-FPS remain less effective at spanning the major spectral directions.

**Figure 11:**
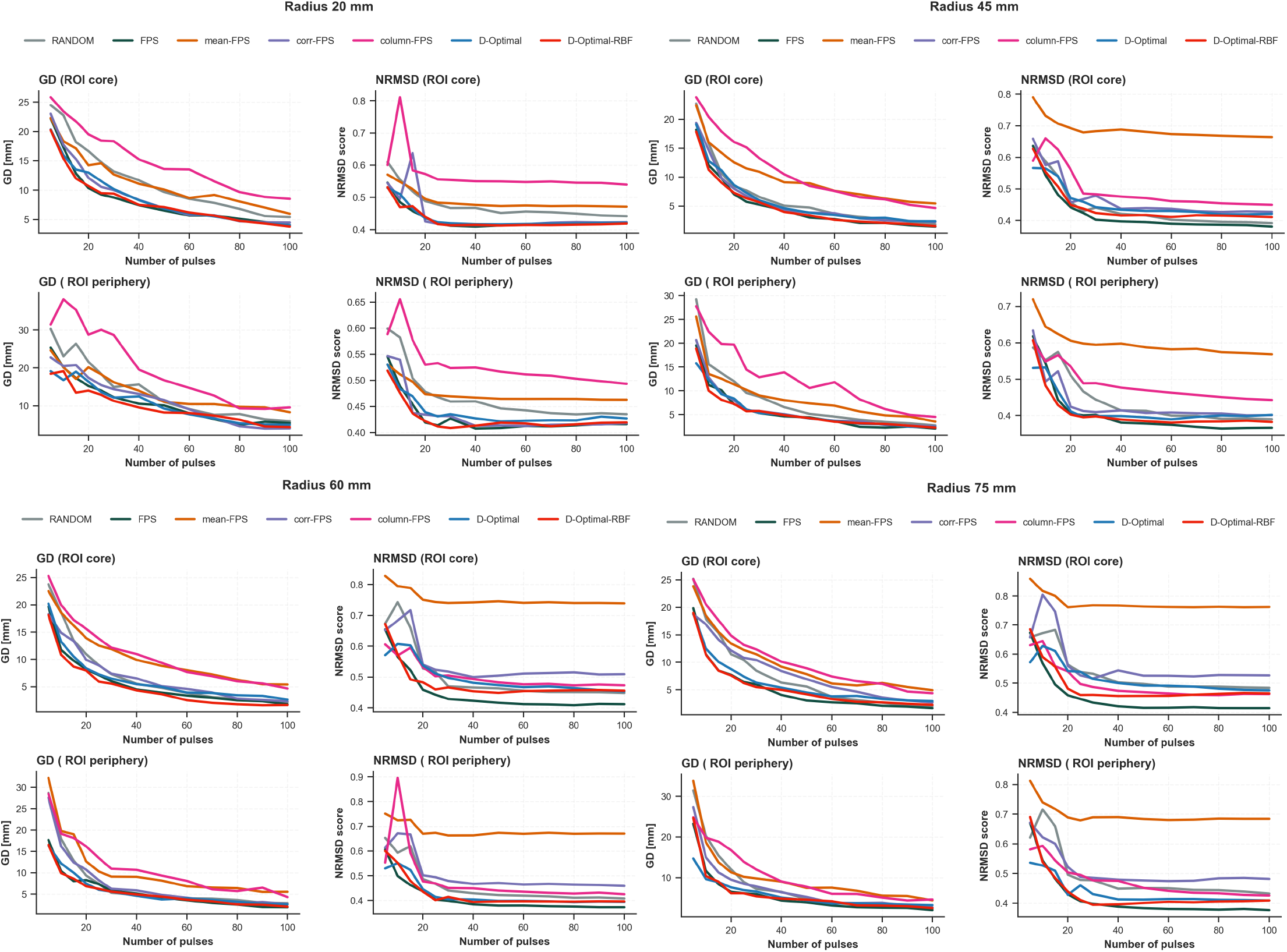
Radius-level robustness across algorithms at broader search radii. Each radius panel overlays all tested algorithms for the 45, 60, and 75 mm search radii.

**Figure 12:**
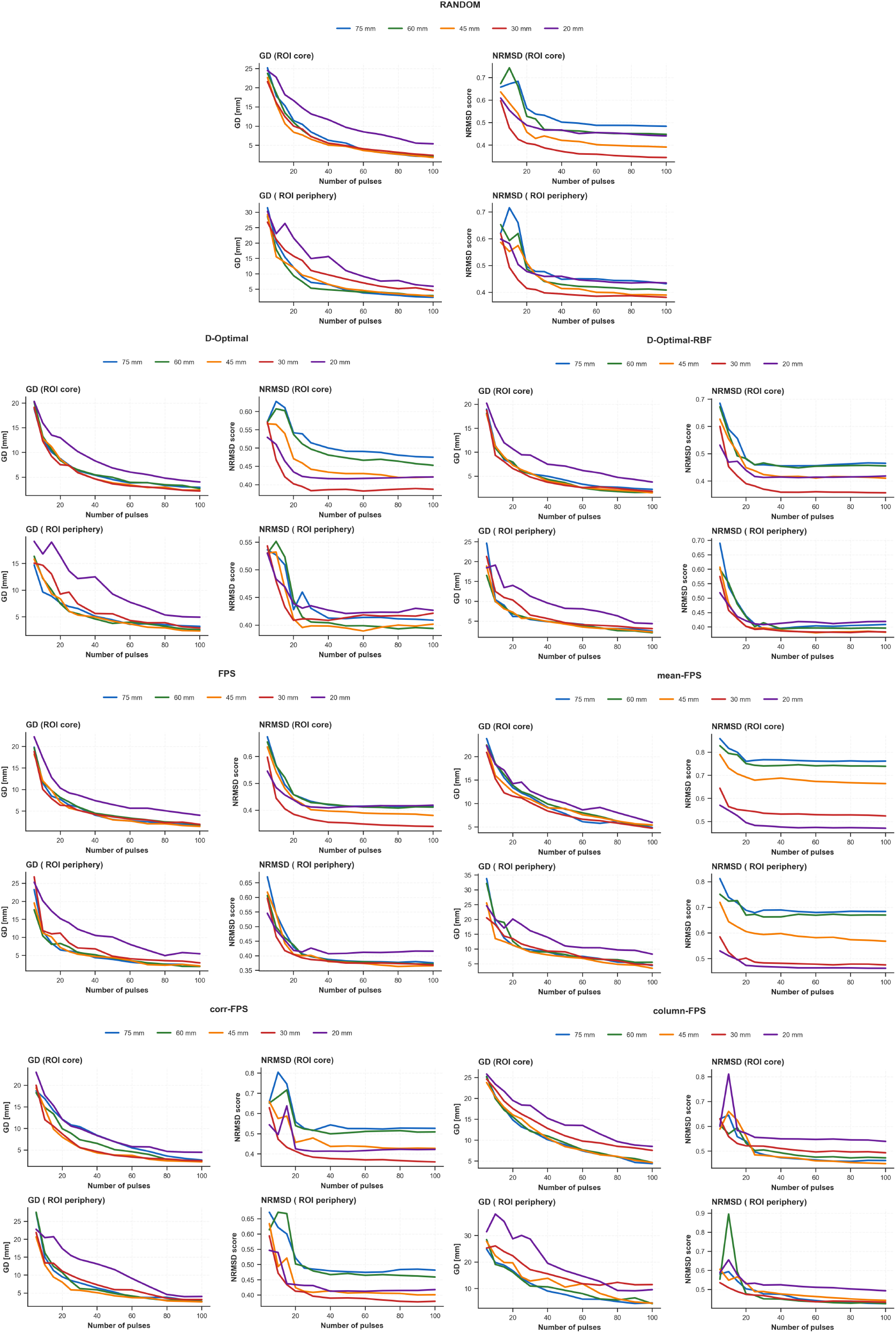
Algorithm-level robustness across search radii. Each algorithm is shown with convergence trajectories overlaid for the 20, 30, 45, 60, and 75 mm search radii. The 20 mm radius produced the most consistent GD degradation across methods and generally worsened NRMSD relative to 30 mm except for mean-FPS, matching the collapsed singular spectrum at the smallest radius. The mean-FPS exception is consistent with its peripheral-attraction bias being partially neutralized when the search region is small.

Mean-FPS improved from 30 to 100 selections but remained less efficient than FPS, corr-FPS, and the D-optimal family, showing that lowering an average correlation-like quantity does not by itself yield efficient subspace filling. Column-FPS had the most negative rank gaps at both budgets, leaving substantial parts of the dominant spectral subspace uncovered. Rank recovery thus tracked method quality but did not fully predict map stability: linear D-optimal led this diagnostic, whereas FPS and kernelized D-optimal were still more stable for later map recovery.

## F Algorithm-level robustness across search radii

The singular-spectrum analysis identified search radius as the only tested discretization hyperparameter that materially changed the candidate E-field library. We therefore tested whether algorithm rankings were preserved across the 20/30/45/60/75 mm radius grid.

Across the tested radii, FPS and both determinant-based objectives preserved their relative rankings, whereas corr-FPS and column-FPS shifted with radius. This asymmetry is consistent with their magnitude handling and with the singular-spectrum analysis: smaller radii do not provide enough E-field diversity, whereas broader radii admit additional weak-magnitude regions, which affects objectives that strip or weaken magnitude information more strongly.

